# Position-dependent variant effects reveal importance of context in genomic regulation

**DOI:** 10.64898/2026.03.17.712488

**Authors:** Sambina Islam Aninta, Ryan Tewhey, Carl G de Boer

## Abstract

Massively Parallel Reporter Assays (MPRAs) provide a high-throughput framework for functionally characterizing how regulatory DNA sequences impact the expression of a model gene. MPRAs have also proven to be useful for measuring the effects of genetic variation, where each allele is typically tested in the center of ∼200 bp of genomic context cloned into the MPRA; but the impact of variant position and local context remains largely unexplored. In this study, we systematically investigate how shifting the position of a variant within an MPRA probe influences its regulatory activity using models that predict expression in MPRAs from DNA sequence. We find that while the direction of variant effects is usually preserved across positions, the magnitude of expression changes can vary substantially depending on where the variant is placed within the construct. This positional bias appears to be largely explained by the strong position-dependent activity of TFs whose binding the variants perturb. In a subset of cases, interactions consistent with cooperativity between TFs also contribute to position-specific effects. ∼1% of variants appear to disrupt RNA polymerase III (Pol III) promoters within *Alu* elements, resulting in position-specificity because both A and B boxes are required for function and exclusion of either motif due to window shifts disrupts the variants’ effects. Consequently, this positional bias can lead to underestimating variant effect sizes from MPRAs, resulting in missed causal variants for downstream functional analyses. Our study demonstrates the complexity of cis-regulatory grammar and how regulatory assays used to study variants can confound their interpretation.

## Background

Gene expression is a product of complex genomic regulation, where TFs bind to specific DNA motifs within cis-regulatory elements such as enhancers and promoters. MPRAs have emerged as a critical tool for studying cis-regulation, enabling detailed dissection of regulatory mechanisms in a controlled context [1–6]. In a typical MPRA design, short genomic sequences encompassing the variant of interest are synthesized and cloned upstream of a minimal promoter and transcription start site (TSS), followed by a reporter gene. Each sequence is tagged with a unique DNA barcode, serving as a sequenceable readout of activity [2]. The library is then transfected or transduced into a relevant cell type where the genomic sequence drives expression of a transcript containing the barcode sequence, and the regulatory activity of the genomic sequence is read out by quantifying the barcodes in the resulting RNA. MPRA datasets have also been used to train deep learning models, which can then be interpreted to reveal regulatory mechanisms [7–9]. For instance, dissection of a model trained on human promoter MPRA data demonstrated that TF binding location can influence regulatory outcomes, with repressor TFs having stronger effects closer to the TSS and activators often having stronger effects upstream [10]. This recapitulated some earlier studies in yeast, which found that most TFs vary in activity by binding location and observed the same trend amongst activators and repressors [11,12].

MPRAs have also emerged as a powerful high-throughput method for functionally characterizing and prioritizing putative causal non-coding variants identified through Genome-wide association studies (GWAS) [3,13–15]. GWAS has identified hundreds of thousands of disease-associated genetic variants, yet most remain functionally uncharacterized, with 90% residing in non-coding regions and confounded with other variants through linkage disequilibrium, making mechanistic interpretation challenging [16,17]. When used for this purpose, each allele for a variant is embedded within the center of a fixed-length window (∼200bp) capturing the endogenous variant context, and tested in the MPRA [8]. Variants are placed in the center to increase the probability that the variant-recognizing TFs are in their proper cis-regulatory context. In addition to testing variant effects, MPRA-trained models are used to predict the regulatory impact of non-coding variants in previously unseen genomic contexts [7,8,10,18,19]. This is a very powerful approach because the number of possibilities that can be explored far exceeds that which can be tested experimentally.

Here, we dissect the regulatory mechanisms that have been learned from MPRA data, and describe their importance for the application of characterizing variant effects. We systematically interrogate how the position of variants within the MPRA probe sequence influences variant effects. We find that the position of variants within the MPRA window impacts predicted variant effects substantially, and that this is supported by experiments testing the same variants in multiple probe locations. Consequently, evaluating the regulatory activity of disease-associated variants should be done with care as their activity may change depending on how the variant is tested. Using computational experiments, we show that these positional biases result from a variety of factors. Firstly, we show that many TFs appear to have differential activity depending on where their motif occurs. Secondly, comparing motif effects in different backgrounds reveals that many TF activities are context-specific. Finally, by testing combinations of mutations, we find that some variants’ effects appear to be mediated by cooperativity-like interactions, with variant effects predicted to depend on secondary motifs. In some cases these correspond to bipartite Pol III promoter elements where the variants’ effects depend on the presence of a motif ∼60 bp away. Overall, these findings support the context-specificity of many TFs and suggest that many disease-causing regulatory variants depend on context-specific regulatory mechanisms.

## Results

### Variant position in the probe sequence affects both predicted and experimentally measured variant effects in MPRA

Given the increasing reliance on MPRA technology to identify causal variants, we sought to determine whether variant effects are modified by location of the variant within the probe sequence. However, evaluating location-specific variant activity at scale requires testing huge numbers of sequences (variants × locations), and so instead we first used a sequence-to-expression model trained on MPRA data to predict the outcome.

To predict variant effects, we employed DREAM-RNN, a state-of-the-art MPRA model, which we trained on a high-quality large-scale human MPRA dataset, generated in the K562 cell line **[7,8]**. The model architecture comprised of a 1-dimensional convolutional layer with a bidirectional LSTM core followed by convolutional layer and global average pooling and was trained for 80 epochs (**Methods**). The model demonstrated strong predictive accuracy for both expression and variant effects, with a held-out test set performance of 0.88 and 0.865 respectively (Pearson’s correlation coefficient) (**Additional file 1: Fig. S1**).

We used this model to predict how variant effects change with position. To test how variant position impacts its predicted effects, we predicted expression changes by systematically shifting each variant’s position across a 200 bp sequence window (**Fig. 1a**). Each window preserves the variant’s endogenous context, but places the variant at a different offset within the reporter construct, so the amount of flanking sequence retained on each side changes with windows. Variant effects are calculated as the difference (ref – alt) in predicted expression (log_2_(RNA/DNA)) between the alleles. We focused our analysis on variants from Open Targets Genetics, which span a range of diseases and traits [20]. In order to enrich for variants that are more likely to be causal for a disease or trait, we only selected single-nucleotide variants from fine-mapped GWAS credible sets where the posterior inclusion probability (PIP) is at least 10%. Most variants in our PIP>10% set are unlikely to alter expression appreciably in K562 cells (for PIP=10%, there is still a 90% chance it is not causal, only some traits are mediated through blood progenitor cells, which K562 is a proxy for, and some variants may not act through altered transcription). Accordingly, we further refined our variant set to include only those variants that showed a predicted variant effect stronger than 0.5 (on a log_2_ scale) at any position. The predicted variant effects revealed substantial positional dependency: the correlation of predicted effects for a given variant decreased steadily as the distance between variants increased (**Fig. 1b**). Similar trends were observed when the log fold change filter was not applied, and all PIP>10% variants were used (**Additional file 1: Fig. S2**). Furthermore, we verified that a model trained on data from a lentiviral MPRA closely mirrored those obtained from the episomal MPRA, suggesting no substantial difference based on the MPRA delivery method (**Additional file 1: Fig. S2**) [9]. Finally, we tested whether the observed positional dependency could be explained by sequence homology between the variants and the training data. Variants with high sequence similarity to the training set were removed using hashFrag [21], and the position dependency analysis was repeated. The positional bias remained among the filtered variants, suggesting that it is robust to potential homology-based data leakage (**Additional file 1: Fig. S2**).

**Figure 1:**
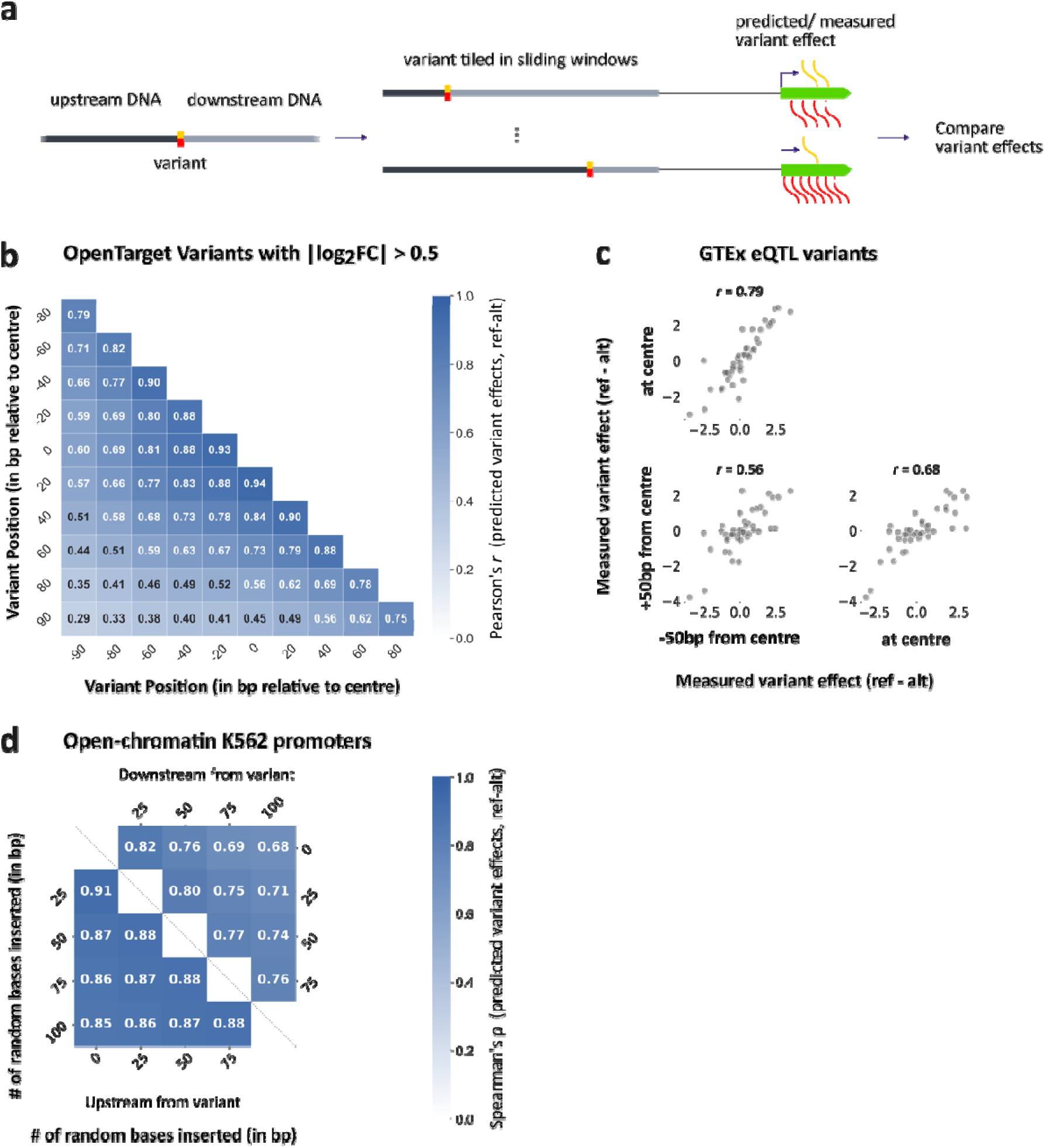
Both measured and predicted variant effects depend on variant position. **(a)** 200 bp probe sequence is extracted such that the variant of interest is tiled at various positions within the sequence window and variant effect (Activity_ref_ – Activity_alt_) is compared to all other windows. **(b)** Correlation of predicted effects (ref – alt) for a given variant decreases steadily as the distance between variants increases. Correlations (Pearson’s *r*; colors) are calculated on the predicted variant effects when variants are placed at different positions (x and y axes). Variants used in this analysis have a PIP>0.1 and predicted variant effect stronger than 0.5 (log_2_ scale) at any position (n=10,862). **(c)** Similar positional dependency is observed in experimental data. Each subplot shows variants (points) with experimentally measured expression changes (x and y axes) when tested at different positions (subplots). **(d)** AlphaGenome predictions of variant effects on RNA-seq in K562 promoters. Inserting random sequence between a variant and the TSS alters predicted variant effects more substantially than inserting random sequence upstream of the variant (so variant remains at the same distance from the TSS). Results across different lengths of random bases inserted (x- and y-axes) are quantified using Spearman’s *ρ* (color).

Upon observing the strong influence of variant position on predicted variant effects, we next sought to determine whether this positional sensitivity could be experimentally validated. We analyzed a subset of MPRA data in K562 cells that tested selected GTEx eQTL variants in three different positions: at the center, 50 bp upstream of center, and 50 bp downstream of center within the MPRA probe sequence [22]. To ensure robustness of the comparison, we only included variants that had a significant variant effect, with an allelic skew Benjamini-Hochberg FDR less than 10% in at least one position, leaving 42 variants. Here too, the closer the variants are, the more correlated their variant effects (**Fig. 1c**). The differences in variant effects was not explained by noise in the data, as we found that inter-replicate correlations were substantially higher than inter-positional correlations (**Additional file 1: Fig. S3**).

### AlphaGenome predicts that variant effects in the genome are position-dependent

Next, we wanted to determine if the position-specific variant effects also occur in the genome. In this case, an equivalent experiment would be inserting DNA of various lengths in endogenous promoters for each of two alleles to change the distance of the variant relative to the TSS and seeing how this changes the native gene’s expression. However, this approach is experimentally infeasible at scale. Consequently, we performed an *in silico* analysis using the state-of-the-art genome-trained AlphaGenome model [23]. For each K562 promoter within open chromatin, we introduced a single synthetic variant 150 bp upstream of the TSS, then inserted 25, 50, 75, or 100 bp of random DNA either 50 bp upstream of the variant (-200 bp from the TSS) or 50 bp downstream, between the variant and the TSS (**Methods**). The predicted effects of each variant (according to AlphaGenome) was then compared with and without each insertion. Expression was quantified at the second exon to ensure that expression changes resulted from altered transcription rates rather than altered TSS usage. In general, inserting DNA between the TSS and variant changed the predicted variant effects substantially more than if the same DNA was inserted upstream of the variant (**Fig. 1d**). Furthermore, more similar length of DNA inserted, more similar distance the variant is from TSS and hence more similar variant effects are observed (**Fig. 1d**), consistent with what was seen in the MPRA (**Fig. 1b, c**). A similar but attenuated trend between insertion length and expression difference was observed when inserting DNA upstream, perhaps as a result of interactions between proteins binding at the variant and those binding upstream of the -200 insertion site. Since inserting additional DNA between the variant and TSS can directly disrupt transcription machinery, downstream insertions have stronger effects than upstream insertions, as expected (**Fig. 1d**). These findings demonstrate that the positional dependence of variant effects is not unique to MPRAs and suggest that the relative position of a variant to other features (e.g. the distance between a variant and the TSS) can change the variant’s impact on expression.

### Potential mechanisms underlying positional dependency of variant effects in MPRA

While our perturbations are unnatural, the positional effects we observe may reflect real biochemical mechanisms underlying endogenous gene regulation. Natural promoters rarely vary element spacing while preserving context, making these context-dependent effects difficult to detect and generalize from genomic data alone. Consequently, we explored the mechanisms that could be at play using a combination of perturbation and prediction with MPRA trained model, DREAM-RNN. We considered some possible mechanisms that could explain how a variant perturbing TF binding could have a positional bias: the TF has position-dependent activity (**Fig. 2a**), and that TF’s activity is modulated by adjacent motifs (**Fig. 2b**) either because the TF is binding cooperatively with another factor in the flanking sequence (**Fig. 2bi**) or because the TF activity itself changes in the presence of other TFs, for instance due to interaction with a co-factor or saturation of a regulatory mechanism (**Fig. 2bii**). Importantly, these possibilities are not mutually exclusive nor exhaustive and multiple such mechanisms could be at play simultaneously in any given sequence. Below, we explore how each of these would manifest.

**Figure 2:**
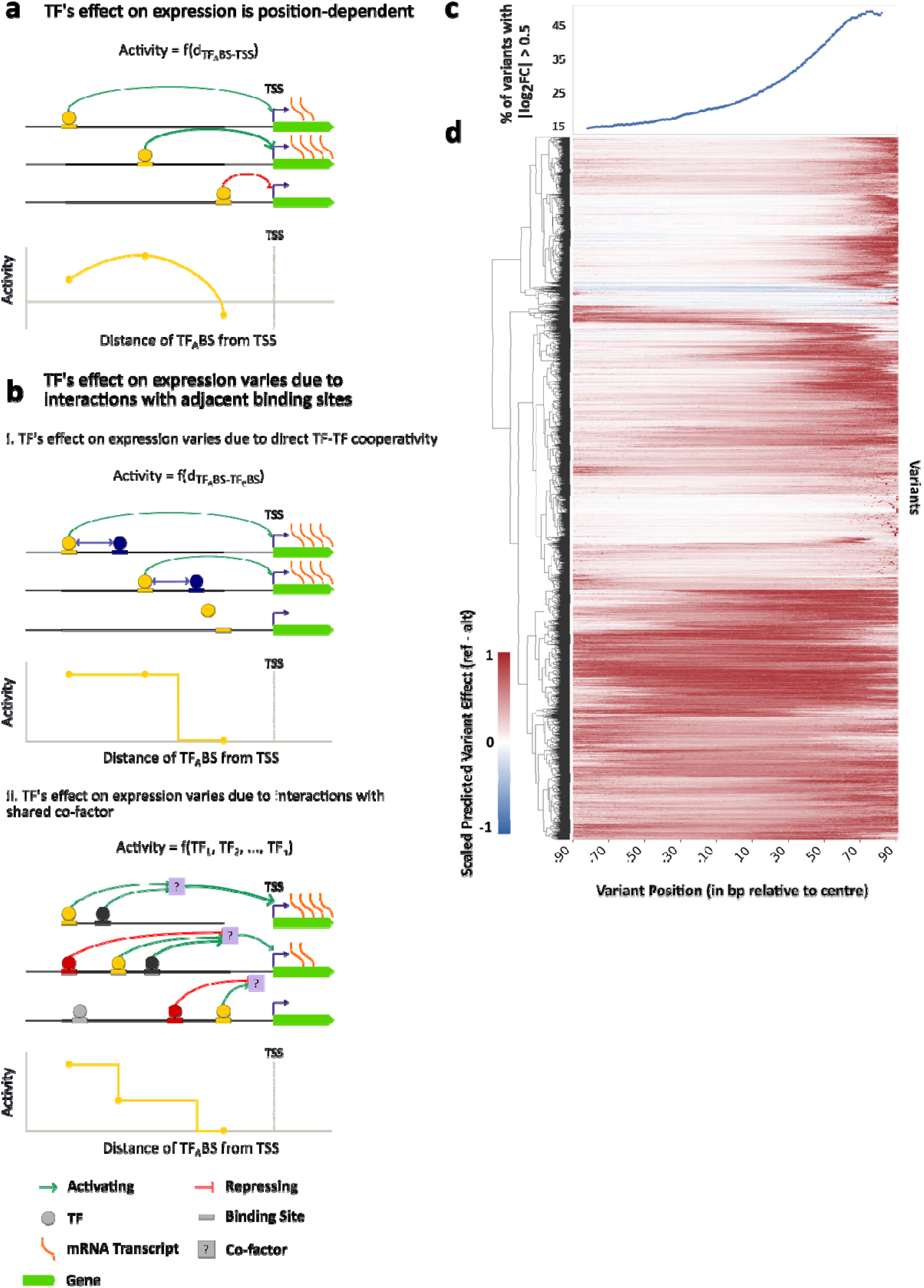
Exploring possible mechanisms explaining positional dependency of variant effects. **(a)** TF’s effect on expression can differ by their distance from the TSS, represented by 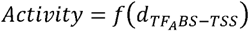 where 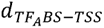 is the distance of the TF binding site from the TSS. **(b)** TF’s effect on expression can differ due to interactions with adjacent binding sites, in two main ways: **(i)** Shifting the variant can create or remove additional TF binding sites that interact physically with the variant-recognizing TF, altering the variant’s effect. This physical interaction means that the TF’s activity is a function of distance between the TF’s binding sites, 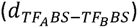 **(ii)** Shifting the variant can create or remove other TF binding sites that modulate the variant’s activity due to interactions with a shared co-factor that differ by variant location. TF’s activity can be defined by *Activity* = *f*(*TF*_1_, *TF*_2_, *TF*_3_, … *TF_n_*) *where TF*_1_, *TF*_2_, *TF*_3_, … *TF_n_* represents all the TFs that interact with a common co-factor. Circles represent TFs, with corresponding binding sites shown as rectangles of the same color. The type and strength of TF activity is indicated by arrow color (green = activating, red = repressing) and thickness. Expression level is indicated by the abundance of wavy lines depicting mRNA transcripts and the gene being transcribed is shown next to the TSS. **(c, d)** Variant effects (ref – alt, predicted using Gosai et al. K562 trained model) vary by position for fine-mapped variants with predicted |log_2_FC| > 0.5 in at least one position (n=33,319). **(c)** More variants have strong effects closer to the TSS. The % of variants with |log_2_FC| > 0.5 (y axis) at each position (x axis). **(d)** Similar positional activities are shared between variants. Variants (y axis) are clustered according to their scaled predicted log_2_FC (colors) for each position (x axis). Each variant is included twice, once in the reverse complement orientation.

Positional variant effects could result from position-specific TF activity. Previous MPRA studies showed that many TFs have position-specific activity that typically varies smoothly across the region tested and sometimes differs substantially by motif orientation [10–12,24–26]. This could result in position-dependent variant effects because the motifs disrupted by the variant could impact expression in different ways depending on where the motifs are located and whether the probe is tested in a forward or reverse complement orientation. In this scenario, the variant impacts the same TF binding site and the TF binding site’s effects typically change only slightly when it is moved over by a single base (but can vary more substantially over long distances). Consequently, we expect that a variant impacting a position-dependent TF binding site will have variant effects that change only slightly as the variant is moved, but can vary more substantially for larger distances or when the motif orientation is flipped (**Fig. 2a**). In this scenario, we also expect that variants impacting the same TF binding site have similar position-dependent expression effects regardless of the flanking sequence context.

Moreover, many TFs are thought to interact cooperatively when binding to DNA, and shifting the variant location could modify cooperative TF binding interactions in the probe by altering which binding sites are present within it [27–29]. If the variant impacts one of a pair of interacting motifs, then the variant effect may depend on whether the interacting site is included in the probe since the two TFs can only interact if both sites are present. How the interacting site influences the variant effect will depend on the nature of the interaction and the variant effect. In some cases, the interaction may be so robust that when both are present the variant has little effect as both TFs remain bound regardless. In other cases, the variant may only have an effect when the interacting site is present as the primary binding site is too weak to be bound in the absence of the interacting site. Regardless, in this scenario, we expect a variant’s effect to be similar regardless of location so long as the two interacting sites are retained in the probe. As the variant’s location is shifted and reaches the threshold at which the interacting binding site is being cut out of the MPRA probe, the interaction can no longer occur, resulting in a sudden change in variant effect; outside this critical threshold location, the variant effect should remain relatively stable since the interacting site is either present, preserving cooperativity, or absent, preventing cooperativity (**Fig. 2bi**). In this scenario, the position dependence would look more like a step function, with the critical position corresponding to when the interacting TF binding site is gained/lost. We also expect that losing the interacting site in other ways (e.g. point mutation) could result in a similar change in variant effect. In this case, we do not expect different variants to have very similar positional dependencies unless they happen to use the same TF-TF pair or a common spacing constraint is present.

Finally, shifting the position of the variant also results in gain or loss of TF binding sites from the flanking region in the probe sequence as genomic sequence context is gained/lost from either end. However, this TF binding site gain/loss would be identical for both alleles. As variant effects are typically expressed as a log fold change between alleles, one would expect that the effects of these secondary binding sites, identical for both alleles, are largely cancelled out. In cases where the flanking sites result in non-additive interactions with the (log) variant effects (e.g. due to interacting with a co-factor or due to approaching saturation), the magnitude of the variant effect could change as flanking binding sites are included or excluded (**Fig. 2bii**). For instance, a variant altering binding of a chromatin opening TF may have an effect only in the absence of other chromatin opening TF binding sites. However, saturating effects should generally not alter the direction of expression change caused by the variant; changes in sign would require more complex biochemical interactions between adjacent factors compared with saturation of a single biochemical function. In this scenario, the changes in TF binding site activity should differ by probe sequence since each would have different flanking TF binding sites. Variant effects may change suddenly as their position is changed due to loss and gain of neighboring sites with which they interact. Since there are not strict spacing and orientation requirements in this scenario, these effects should vary by variant since neighboring TF binding sites will differ in position and identity for all variants.

### Causal variants might be underprioritized depending on their position within the probe sequence

To test our proposed hypotheses, we first sought out to determine if there were common trends in how variant effects changed with respect to position. We predicted variant effects of the Open Target variants (following the same approach as in **Fig. 1a**) at 1 bp resolution and clustered the variants based on the Euclidean distance between predicted effects (**Fig. 2c, d**). Our results show distinct clusters (**Fig. 2d**). The vast majority of variants are consistent in direction of change across positions, but there are a few variants that change signs. Most variant effects tend to be smooth, with a single variant effect maximum, inconsistent with a context dependency where a variant effect might increase or decrease as nearby sites are gained and lost during variant shifting (**Fig. 2b**). However, this is consistent with what would be expected if the variant-affected TFs had position-specific activity (**Fig. 2a**), which has previously been observed to be smooth and without frequent sign changes [11]. About half of variants have a reasonably consistent effect across most of the probe region, with a maximal effect at varied locations within the probe, most commonly proximal to the TSS (**Fig. 2c, d**). Another sizable group of variants have relatively weak effects except for one location, usually proximally to the TSS. Within this group, there are some where the large variant effect is restricted to a single or very few positions, suggestive of an artifact resulting from interactions with the constant flanking sequence context of the MPRA (e.g. a TF binding site is formed that bridges the MPRA probe and constant flank). Moreover, the precipitous changes in activity that would be most consistent with gain or loss of a cooperative interacting binding site (**Fig. 2bi**) do not appear to be common (**Fig. 2d**), but we expect that these would not form a single cluster as the locations of the precipitous changes in variant effects would likely be different for every variant, depending on type of cooperative interaction and the location of the interacting site. Overall, the proportion of variants with |log_2_FC| > 0.5 increased closer to the TSS, plateauing at very close distances (**Fig. 2c**). In the context of testing variant effects to prioritize trait-associated variants, the variants’ placement within the probe will impact their effect sizes in the MPRA, potentially misclassifying true regulatory variants as having minimal effect.

We repeated this analysis using a model trained on lentiMPRA data from Agarwal et al. to ensure that our results are robust to MPRA delivery method and found very similar results (**Additional file 1: Fig. S4**) [9].

### TFs show position-dependent activity for human and yeast

We next further tested the hypothesis that TF activities varying by position could help explain the positional dependency of variant effects (**Fig. 2a**). We restricted our analysis to the known critical TFs in K562 [30–32], expecting that positional effects for these TFs should be most readily detected. The consensus sequence of each motif was embedded into 500 random DNA probes of 200 bp in 1 bp increments (**Fig. 3a**). We then performed complete *in silico* knockout of the motifs and averaged this knockout effect (WT – KO) across the 500 random sequences to assess how disruption of TF motifs in different positions affect overall contribution to regulatory activity. Our analysis revealed diverse positional effect patterns across different TF motifs, with most having strongest effects closest to the TSS (**Fig. 3b**), similar to what was seen with variant effect predictions (**Fig. 2d**). Additionally, for a few TFs, motif orientation also played a role in this positional effect, where the regulatory effect is a stronger or different in one orientation, suggesting a strand bias in TF activity (e.g. EWSR1). The majority of TF motifs had a consistent direction of effect on expression (e.g. activating or repressing) regardless of placement, but several (e.g. FOX heterodimeric motifs, UME6) were not, similar to what was observed with variant effect predictions (**Fig. 1c**, **Fig. 2d**). Some motifs (like NRF1 and YY1) showed weaker positional effects than were reported before [10]. One likely explanation is type of assay: in the previous literature, positional effects were shown using a promoter MPRA assay, whereas the model we used here to predict TF’s position dependency is trained on proximal enhancer assay data, with sequences tested upstream of a fixed core promoter. Since YY1 primarily binds near the TSSs, and the MPRAs only tested sequences upstream of the TSS, our MPRA models did not learn TSS-proximal effects.

**Figure 3:**
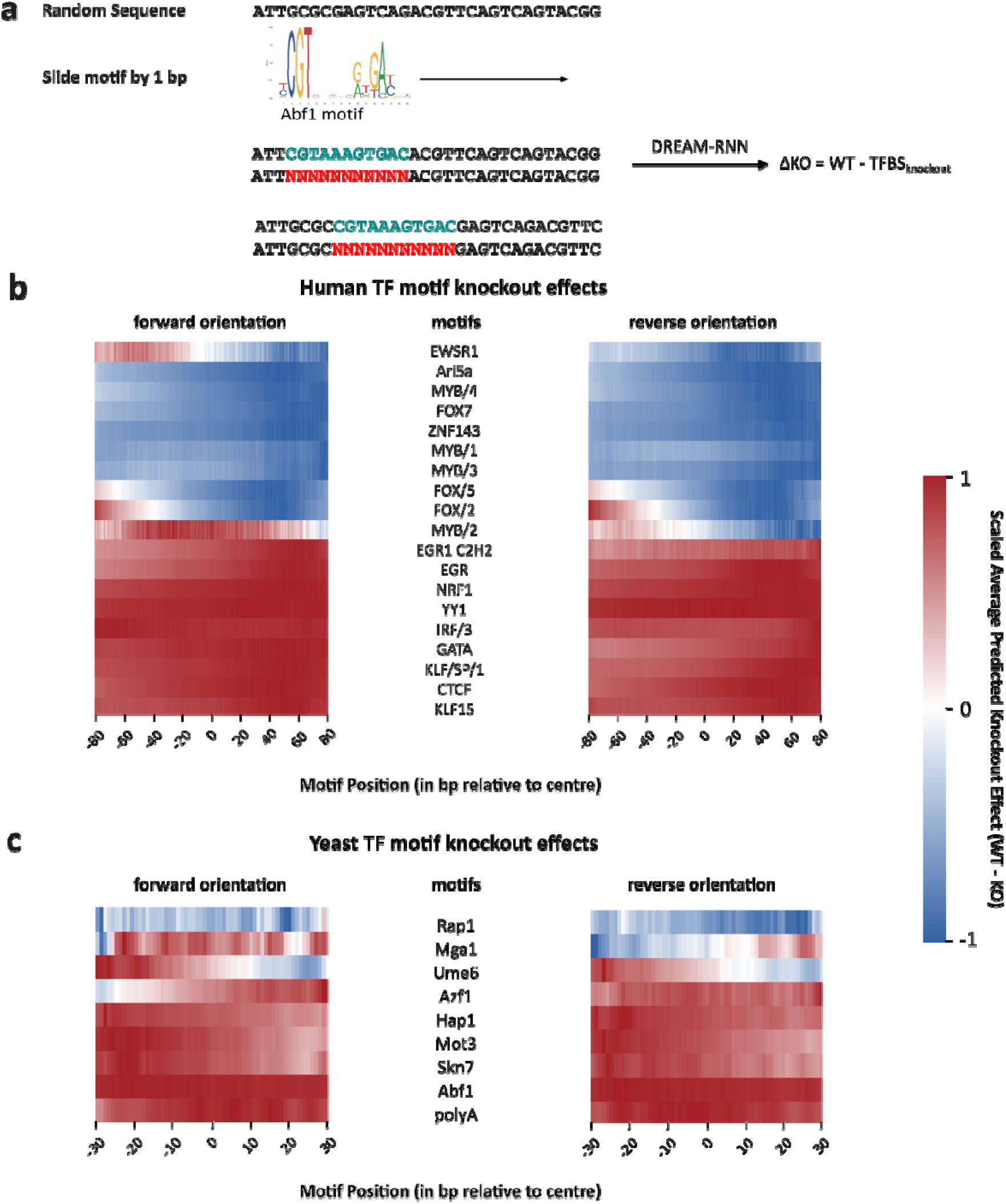
TFs exhibit position and orientation-dependent activity in both yeast and human. **(a)** The motif of interest is tiled every 1 bp across moderately expressed random sequences and the motif knockout effect (predicted change in expression when the motif is replaced with Ns) is averaged across all random sequences for each motif. Each motif is tested in the forward and reverse complemented direction. **(b-c)** Scaled and averaged predicted knockout effect (WT – KO) (colors) for each TF (y axes) across multiple positions (x axes) for **(b)** human and **(c)** yeast. Knockout effects here are scaled by the maximum absolute effect (strongest effect regardless of sign) observed at any position within each TF.

As a control, we performed the same analysis using yeast random promoter-trained DREAM-RNN model, where we know from previous analyses of the same data that many motifs are strongly positionally dependent [7,11]. In particular, we compared the positional biases for each TF learned by a physics-informed neural network to our *in silico* knockout of motifs [11]. Positional and orientation dependencies were evident, including the previously observed ∼10.5 bp periodic biases consistent with the helical periodicity of DNA (**Fig. 3c, Additional file 1: Fig. S5**).

Overall, our results from both human and yeast (**Fig. 3b,c**) support the hypothesis that TF activities vary by position (**Fig. 2a**) and explain why variants in these TFs could have different regulatory effects depending on their positions.

### Probe sequence context alters the knockout effect of TFs

Next, we asked whether the surrounding sequence context within the probe could alter the effect of a TF motif. It has been previously shown, through *in silico* experiments, that the surrounding context can modulate TF activity [18]. Such changes could also arise when flanking sites introduce non-additive effects: for example, if additional flanking sequences contain motifs that are redundant with the first (**Fig. 2b**). To test the context dependence of TF motifs, we embedded human motifs for TFs that are active in K562 cells into both random DNA sequences and genomic sequences (putative K562 enhancers from Gosai et al. [8]), placing the motif at the center of the probes, and compared the predicted expression of these to matched sequences where the motifs were replaced with Ns (**Methods**). We expect that the effect of the TF on expression will be reflected by the change in predicted expression observed when its cognate motif is included; any context specific effects will be evident in the distribution of expression changes across many sequence contexts. However, we also note that other possibilities can impact the motif effects as tested in this way, such as secondary motifs occurring that overlap the one inserted and are recognized by other factors. Accordingly, we also include random control motifs – shuffled versions of the originals – to illustrate the distributions expected by chance (**Methods**).

The random DNA and putative enhancer backgrounds largely showed similar trends for the same TF. In both, the predicted knockout effects (WT – KO) for each motif varied substantially between sequences into which the motifs were embedded (**Fig. 4**). Compared to the random control motifs, most motifs we tested were activating. CTCF and KLF13 motifs were activating in the vast majority of sequences. Several others were usually activating but nonetheless had a repressing effect in many contexts (e.g. EGR1, IRF5). The remaining motifs (e.g. EWSR1, FOX/5) frequently differed in the direction of knockout effect between embedded sequences. While there does appear to be context dependency of TF activity, it is largely inconsistent with saturation effects, where we would expect to see TF motif knockout effects plateau at 0 (e.g. where the TF binding site becomes wholly redundant); this kind of plateau was observed only for a few of the motifs such as CTCF, YY1 (**Fig. 4**). However, motifs will sometimes have opposing effects due to fortuitous creation of binding motifs for other TFs that overlap the motif being tested, so we may not expect a strict plateau at 0 even with saturation. Nevertheless, the fact that many of the TFs tested here have a wide distribution of effects when their motifs are fully deleted suggests that single base changes to those same motifs could also have variable effects on expression, in support of the context-dependence hypothesis (**Fig. 2b**).

**Figure 4:**
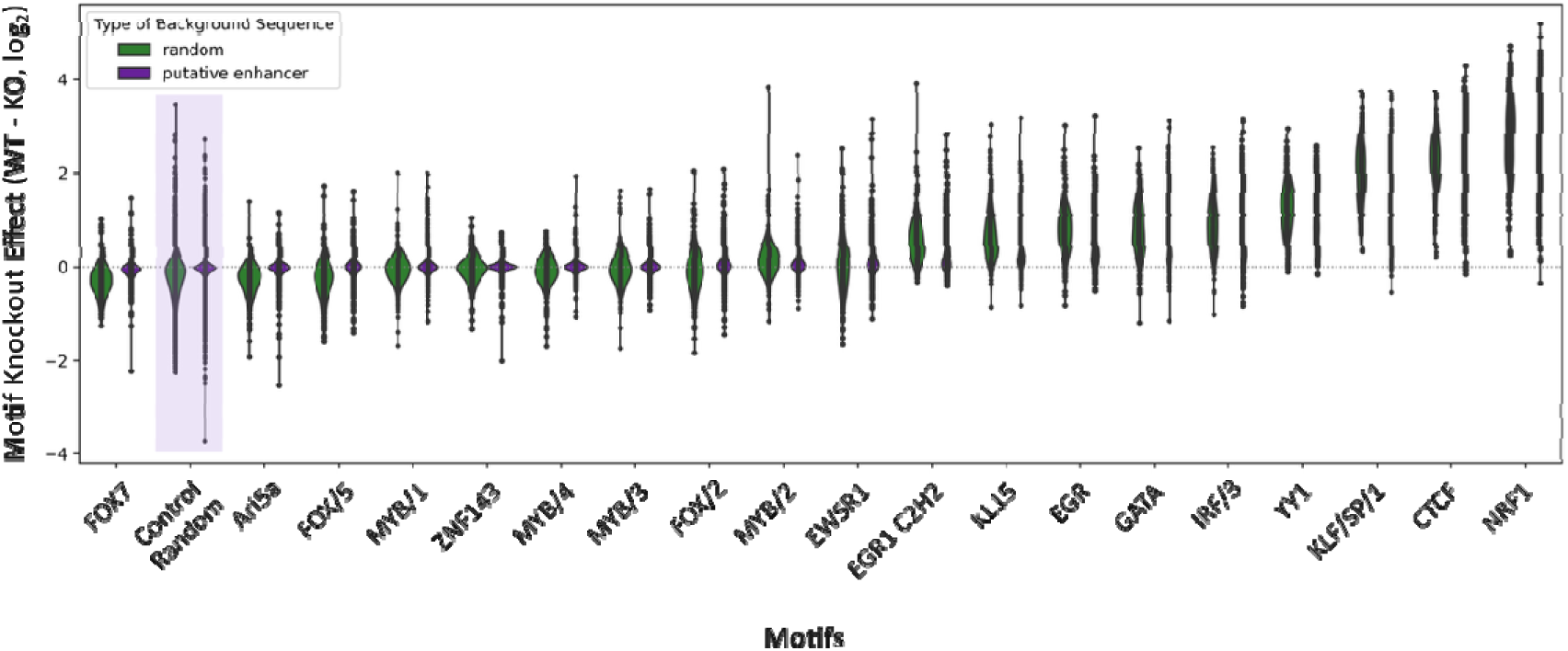
Flanking probe sequences influence the predicted knockout effects of TF motifs. Distribution of predicted motif knockout effects (WT – KO) (log_2_ scale) (y axis) when the motif (x axis) is placed in the center of random (green) or putative genomic enhancer (purple) sequences. Random sequences (n=20) were included as controls and are highlighted in the figure for comparison with the motifs. The violin shape represents a kernel-density estimate of the distribution, with width corresponding to value frequency, and the central point marking the median.

### Epistasis between variant and distant neighboring motif consistent with the bipartite RNA pol III promoter architecture

We next sought to analyze the mechanisms of differentially active variants in more detail. We selected the experimentally determined variant, rs2316952, (**Fig. 1c**) that showed the largest change in both predicted and measured effect across the three positions. We identified the key nucleotides contributing to the model’s predictions for each allele and position via *in silico* mutagenesis (ISM), effectively testing for epistasis between the variant and other potential mutations in the sequence (**Methods**). Consistent with the measured position-dependence of this variant, the motif at the site of the variant only appears to be important in two of the three tested positions (**Fig. 5a**). There was a second important motif (box B) about 65 bp away from the variant-affected motif (box A) that is much more important when the high-expression allele was present. When box B is lost due to the positional shift, the variant is predicted to have little impact and the box A was no longer important, suggesting the box A depends on box B. In contrast, when the motifs were shifted but both retained within the sequence, the variant effects were similar and both motifs remained important to the model’s predictions.

**Figure 5:**
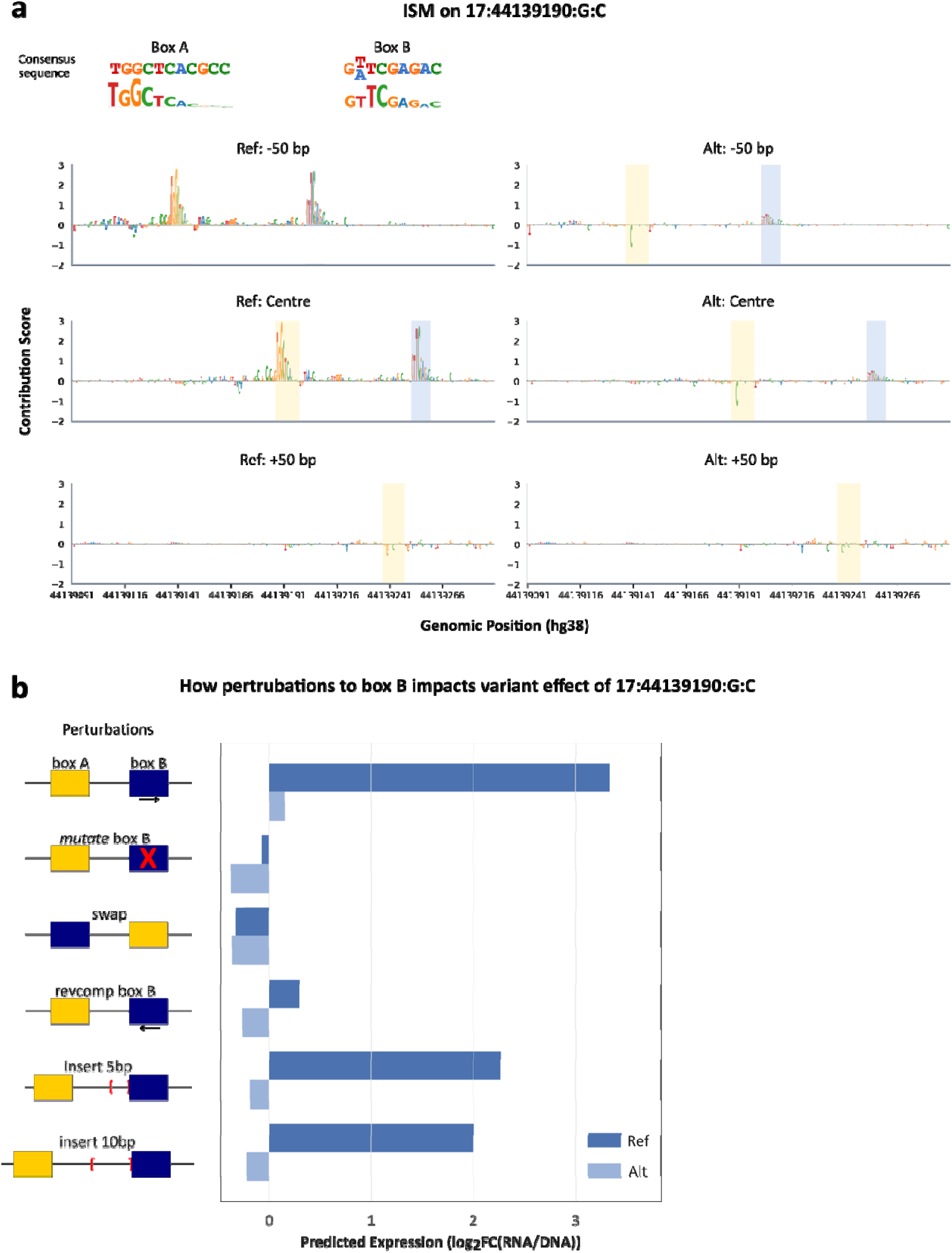
Shifting a variant can alter its activity by partial loss of pol III promoter elements. **(a)** Average contribution scores (y axes) across 200 bp probes (x axes) with the variant at three prediction windows (rows). The two most important regions, labeled as box A and box B (colors), closely match the known A and B box consensus sequences of the Pol III promoter (top) [34]. **(b)** Perturbations to A and B box interaction and predicted impacts on expression. For each perturbation (rows; legend far left), we predicted expression (x axis) for Ref and Alt (colors).

Upon further investigation and alignments with RepeatMasker, we found that the box A and B sequences have high sequence homology with the type 2 Pol III promoter found in *Alu* elements (**Fig. 5a**) [33–35]. Type 2 Pol III promoters are internal promoters: box A and box B elements located within the transcribed region recruit TFIIIC and TFIIIB directly, allowing Pol III to initiate transcription without upstream regulatory sequences [36–38]. *Alu* elements on the other hand are SINEs that exploit this architecture: they contain embedded Type 2 Pol III promoter elements (box A and box B) within their sequence, enabling Pol III-mediated transcription and retrotransposition, hijacking the cell’s own transcription machinery. *Alu* variants have previously been tested in an episomal MPRA where Pol III promoter regions were observed to drive regulatory activity [35]. Since the K562 MPRA training data from Gosai et al. was not derived from a poly-A capture assay, it likely also captured Pol III regulatory activity, which the model had learned and subsequently reflected in the importance assigned to box A and B motifs at this variant [8].

We wanted to test what aspects of Pol III regulation our model has learned and gauged the impact of various mutations and rearrangements of these motifs (**Fig. 5b**). To transcribe *Alus*, TFIIIC first recognizes box B with high affinity, which anchors the complex to the *Alu* element, and then contacts box A upstream to complete assembly and recruit TFIIIB and subsequently Pol III [39]. Consistent with this cooperative bipartite recognition mechanism, mutating box B had a similar effect on predicted expression as when box A is disrupted by a variant (**Fig. 5b**). Similarly, reverse complementing box B or swapping the locations of the two motifs were both predicted to reduce expression and diminished the importance assigned to box A (**Fig. 5b**), consistent with the strict orientation requirement of the Pol III promoter where box A has to be upstream of box B relative to TSS for proper TFIIIC function [39]. Moreover, inserting random bases between the two motifs had minimal effect on predicted expression, whether for 5 or 10 bp (∼0.5 and ∼1 DNA helical turn) (**Fig. 5b**). This is consistent with the known architecture of TFIIIC, in which the τA and τB subcomplexes that recognize box A and box B, respectively, are connected by a flexible linker that can accommodate variable spacing. Together, these observations are consistent with the known bipartite architecture of Pol III promoters in *Alu* elements, and therefore we concluded that the model has learned characteristics of Pol III regulatory activity.

Furthermore, this finding provides a concrete example of how position-dependent differences in TF cooperativity can influence variant effects, supporting our hypothesis from **Fig. 2b(i)** where losing box B loses the cooperative interaction so the variant in box A no longer causes a change.

### The majority of cooperativity-like distant epistasis corresponds to Pol III promoter elements

While the previous example (**Fig. 5**) reflected the expected epistatic interaction between the two core motifs of an RNA pol III promoter, cooperativity between TFs would be expected to have similar effects, and so we next sought to determine how frequently such interactions contribute to variant effects. To investigate this, we analyzed Open Target variants with strong predicted effects when the variant was placed at the center of the probe (|log_2_FC| > 0.5). For each variant, we performed ISM across all positions for both the high-expressing allele and low-expressing allele. To assess how per-position importance differs between the two alleles, we then computed the difference in the ISM score (**Methods**). This captures the epistasis between the variant and each of the other positions in the sequence, indicating some interaction.

We found that epistatic interactions of differing strength between the variant and other positions in the sequence (**Fig. 6; Additional file 1: Fig. S6**). We expect epistasis in the vicinity of the variant to be common because mutating the same motif in multiple ways will produce non-additive (epistatic) effects on expression. Given the size of a typical motif, the epistasis resulting from a single motif being perturbed is likely restricted to about 21 bp region surrounding the variant (∼10 bp to either side). However, direct cooperativity between TFs often forms a compact compound motif, shorter than the sum of the two individual motif lengths [40], and so will likely operate within a range that is difficult to distinguish from single TFs. The majority of variants did not form obvious clusters, indicating a lack of a common mechanism (**Fig. 6a**). We investigated the variants that clustered with similar long-range interactions (**Fig. 6a**), which made up ∼1% of the total Open Targets variants with |log2FC| > 0.5, and found that they closely matched *Alu* elements and were specifically variants perturbing the box B motif, part of the bipartite Pol III promoters **(Additional file 1: Fig. S7)** [34].

**Figure 6:**
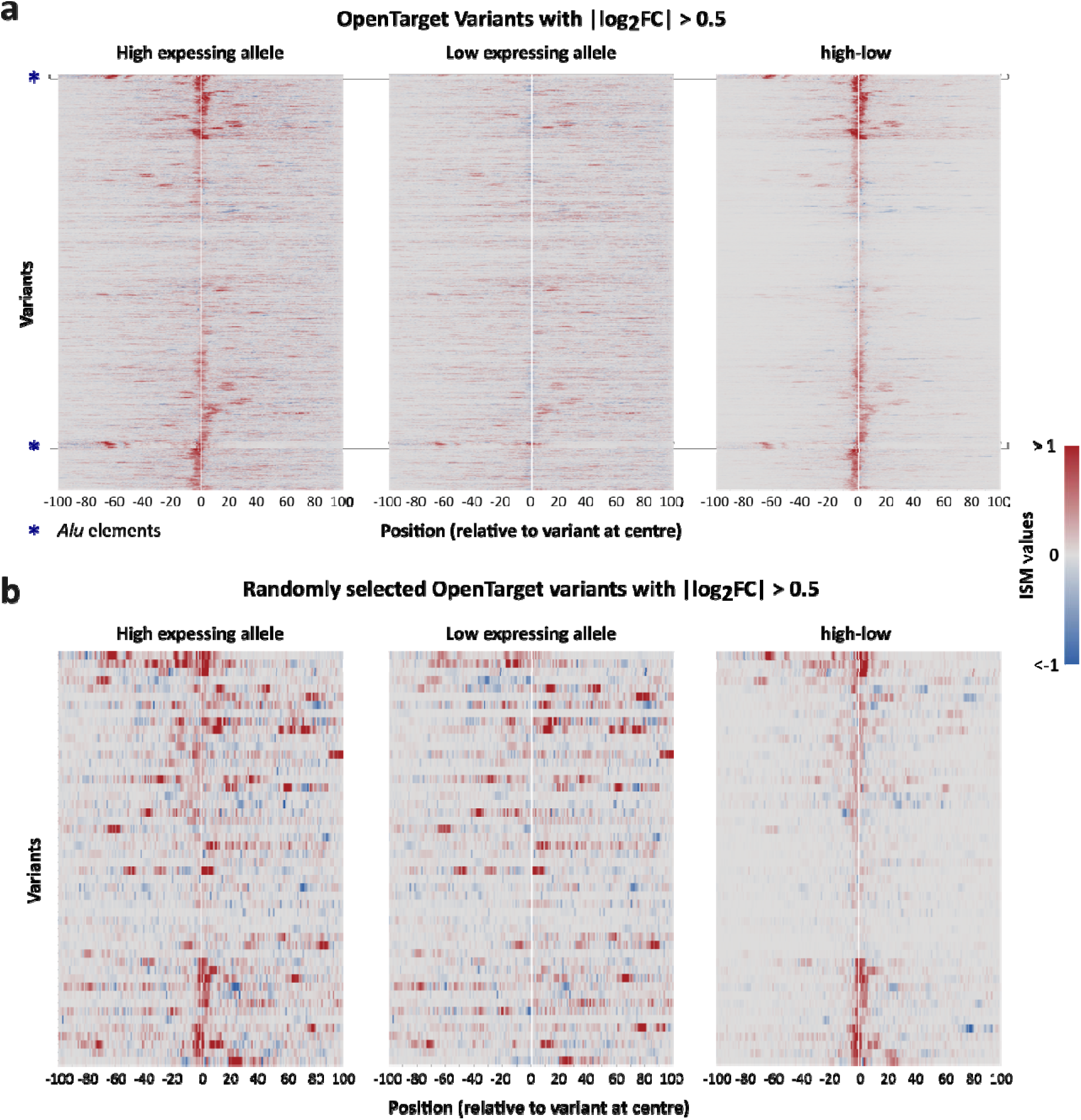
Predicted epistatic effects in OpenTargets variants suggests cooperative interactions modulating variant effects are not common. **(a)** Each row represents a variant with |log₂FC| > 0.5 (n=2956). The *Alu* elements exhibited strong long-range interactions. **(b)** Randomly selected 50 variants from the 2956 variants in (a) to show typical interactions. The heatmaps show average ISM contribution scores (color) across 200 bp sequences (x axis), where variants are centered. The high-low panel shows the difference in ISM values between the higher- and lower-expressing allele profiles. Variants are clustered based on the Euclidean distance of high-low.

## Discussion

MPRAs have proven to be a promising approach for measuring variant effects because of their high-throughput and ability to enrich for causal variants [3,15]. However, our analysis shows that variant effects measured using MPRA can be highly positional and strand specific, potentially resulting in mis-prioritization of variants. At present, MPRA probe sequences are typically around 200 bp, and ongoing improvements in oligo pool synthesis enable close to ∼400 bp at present. Longer probes could help preserve more surrounding regulatory information in case it depends on complex interactions with other factors. However, longer oligos come with increased error rates in probe synthesis, which can impact probe expression and potentially obscure true variant effects. In addition, nucleosome positioning at the endogenous locus is likely to differ from that in MPRA constructs, and shifts in nucleosome placement can drastically change regulation by exposing some motifs and masking others. Designing MPRA probes to align with the edges of open chromatin regions, rather than arbitrarily centering variants within synthetic sequences, could more accurately reflect the native genomic environment. On the other hand, we also observed that the direction of a variant’s effect was usually consistent across the probe and generally easier to detect closer to the TSS for most but not all variants (**Fig. 2c; Additional file 1: Fig. S4**), suggesting that longer probes (where the variants are even further from the TSS) may not help in most cases. Testing each variant in several locations may maximize the chances of seeing an appreciable change. An alternative is to use deep learning models trained on the MPRA data. These models will have learned these positional biases, and it is possible that a sufficiently good model could be used to estimate positional biases in variant effects and potentially compensate for them to get a better estimate of the endogenous variant effect. It may soon be more efficient to perform an MPRA designed to generate the best training data and then predict variant effects, rather than measuring the variants’ effects in an MPRA directly, since the model can be used both to dissect variants and their regulation in ways that are not feasible to test in an MPRA.

Our results suggest that the positional dependencies observed in these MPRAs reflect fundamental regulatory constraints at endogenous genes. Motif occurrence across human CREs is highly non-random; approximately 35% of human TF motifs are biased in their occurrence by location within enhancers as well as promoters, clustering within specific territories relative to the TSS [30,41]. While we found TF motifs have a position and context dependence (**Fig. 3, 4**), this was itself in the context of an MPRA system; we expect that similar rules will be at play in natural promoters, but their exact manifestations will, of course, depend on each promoter’s context (e.g. what binds there and where).

Furthermore, we identified several variants located within RNA Pol III promoters embedded in *Alu* elements (**Fig. 6**). In these cases, two spatially separated interacting regions were detected, which aligned with the bipartite promoter architecture defined by box A and box B motifs (**Fig. 5a**) [33,35]. This observation is particularly interesting, as Pol III transcription requires the coordinated activity of both promoter elements. Disruption of either motif, failure to include both motifs within the tested window, or testing the promoter in the wrong orientation would affect promoter function. These findings therefore highlight the importance of sequence context in variant interpretation and shows how accurate prioritization of Pol III variants depend on ensuring that both boxes are tested in MPRA in the right orientation.

Finally, while we saw little evidence that the positional dependence of variant effects resulted from redundancy between motifs (**Fig. 2bii**), we did see that the activity of TFs appeared to be highly context dependent (**Fig. 3**, **Fig. 4**), suggesting that the interactions with nearby bound factors can influence the activity of a TF motif (and, consequently, variation in that motif). This context dependence of TF motifs can be challenging for deep neural networks to learn, particularly when training data are limited, and may help explain why state-of-the-art sequence-to-function neural networks like Enformer and AlphaGenome still struggle to consistently predict the correct direction of variant effects [23,42,43]. Nonetheless, our AlphaGenome analysis suggests that it has learned at least some of the context dependence of TF motifs.

While we did validate the positional-dependence of variant effects with experimental data (**Fig. 1c**), the vast majority of our analyses are based on computational predictions of variant effects. As the scale and sensitivity of MPRAs increase, these predictions could be experimentally verified in more detail to better understand the mechanisms at play.

## Conclusions

In summary, we show that the position of a variant within an MPRA probe can substantially change its measured effect, mainly because TF activity depends on position and sequence context. Applied to fine-mapped variants from GWAS, our analyses reveal that this positional bias can lead to underestimated effect sizes and missed causal variants. Therefore, our work highlights the context dependence of cis-regulatory activity and explains how using MPRA to prioritize non-coding variants could underprioritize causal variants.

## Methods

### Selecting Open Target Variants

GWAS SNVs with PIP at least 0.1 from the Open Targets dataset were selected, resulting in 103,353 fine-mapped variants spanning a range of diseases and traits [20,44]. For each variant, 200 bp windows containing the reference and alternate allele were extracted from the hg38 genome, with the position of the variant within the window varying by experiment (**Fig. 1a**).

### DREAM-RNN overview and training

The DREAM-RNN model was trained on K562 MPRA data following the procedures described in Rafi et al. [7]. This model was selected for its state-of-the-art performance in predicting expression from MPRA data. The architecture combines 1D convolutional layers (kernel sizes 9 and 15 bp, capturing sequence motifs) with a bidirectional LSTM core (640-dim hidden states) followed by convolutional layer and global average pooling for expression output. Training was performed for 80 epochs with a learning rate of 0.005, using 230 bp input sequences in which the 200 bp variable region was flanked by 15 bp adapter sequences on each side. The model outputs predicted MPRA expression, as log_2_FC (RNA/DNA) (the RNA produced divided by its abundance in the DNA library). To account for differences in MPRA delivery methods, separate models were trained on K562 MPRA datasets from Gosai et al. and Agarwal et al., corresponding to episomal MPRA and lentiMPRA respectively [8,9]. Unless otherwise stated, we used the Gosai-trained model for all analyses.

### Analysing positional dependency on predicted variant effects

For prediction, each 200 bp sequence containing an Open Targets variant was padded with 15 bp adapter regions, using adapter sequence from either the Gosai MPRA or Agarwal MPRA depending on the model applied. Using the trained K562 DREAM-RNN models, expression levels (log_2_(RNA/DNA)) for the reference and alternate alleles were predicted, and variant effects (reference – alternate) were calculated at each position. To test whether sequence orientation affected the results, each 200 bp sequence was reverse complemented and variant effects were predicted in the same way (keeping the 15 bp adapter same as the forward orientation). Since many variants are unlikely to have measurable effects in K562, only those with at least a variant effect of 0.5 (log_2_) in at least one position were retained for downstream analyses unless otherwise noted.

The variant effects at different positions were further analyzed to assess positional dependence. First, we calculated the Pearson’s *r* to evaluate how well variant effects at each position correlated with those at other positions (**Fig. 1b**). Next, the variants were clustered using Euclidean distance between their variant effects (**Fig. 2d**). Since the reference and alternate allele assignments are arbitrary with respect to the variant’s effect, each variant effect was scaled by its largest absolute effect across positions, effectively normalizing all variants to have a maximum effect of +1. Both forward and reverse complement sequences were combined during clustering. For clustering analysis, 33,319 variants were selected from the predictions made at 1 bp.

To evaluate whether the delivery method of the reporter construct library influences positional dependency, both the clustering and correlation analyses were repeated using models trained on K562 data from Agarwal et al. and Gosai et al. (**Additional file 1: Figs. S2 and S4**).

Finally, to evaluate how homology-based leakage with the training data affects the model-predicted positional dependency, the variants were filtered using hashFrag based on their sequence similarity to the training set using default parameters [21]. The clustering and correlation analyses were then repeated on this subset of variants using models trained on the K562 data from Gosai et al. (**Additional file 1: Figs. S2 and S4**).

### Experimental Variant Effect Analysis

GTEx variant MPRA data were obtained from Siraj et al. [22]. Variants were first filtered to retain only those with data available across three positions: variant placed at center, variant at 50 bp upstream and 50 bp downstream from center. Variants were then further filtered to ensure that both reference and alternate alleles were present. To make sure these results did not result simply from noisy measurements of variant effects, only variants with a significant variant effect were selected. Welch’s *t*-tests were performed between the reference and alternate alleles using the five replicates per allele, without assuming equal variance. Multiple testing correction was applied using the Benjamini-Hochberg false discovery rate (FDR), and variants with FDR < 10% in any of the three positions were retained. Pearson’s *r* was then calculated between each pair of positions (**Fig. 1c**).

Inter-replicate correlation of variant effects was assessed by bootstrapping the five replicates. Replicates were resampled without replacement 10 times (number of unique ways to divide five replicates into groups of two) and split into two groups, and Pearson’s *r* was computed between variant effects across the two groups, yielding a distribution of inter-replicate correlations (**Additional file 1: Fig. S3**).

### Predicting K562 promoter variant effect using AlphaGenome

We identified known K562 promoters located within open chromatin regions and retained those with sufficient flanking sequence such that exon 2 was fully contained within a 131,072 bp sequence window centered on the TSS (n=17,535). For each promoter, we introduced a single transition variant (A↔T or C↔G) at 150 bp upstream from TSS and inserted random DNA sequences of 25, 50, 75, or 100 bp either (1) upstream of the variant, placing the variant 200 bp upstream of the TSS, or (2) between the variant and the TSS, placing the variant 100 bp upstream of the TSS. To maintain a constant input length, an equivalent number of bases was removed from the far upstream end of the sequence, ensuring that the TSS and exon 2 locations stayed fixed within the prediction window. Reference and alternate sequences for each condition were then evaluated using the pretrained AlphaGenome model (K562 RNA-seq track from PyTorch reimplemented port was used [45]). Variant effects were quantified as the predicted difference in expression between the reference and alternate alleles, measured as the mean across exon 2, which was used in preference to exon 1 because exon 1 frequently contains untranslated regions (UTRs) and our perturbations could have interfered with the predicted TSS location. For each pair of perturbation types, predicted effects for the same variants were compared with Spearman’s *ρ* (since Pearson’s *r* is more affected by outliers).

### Analysing position specific TF motif activity

Consensus motif sequences of the TFs active in K562 and yeast were obtained from Vierstra et al. and de Boer et al. respectively [11,31]. The motifs were embedded in a background of random sequence, in sliding windows of 1 bp across the random sequence. *In silico* knockout effects (replacing the motifs with Ns by setting each base at that position to 0.25 in the one hot encoding) was performed with the motifs in different positions and predicted using DREAM-RNN. Knockout effects for each position for each motif were averaged across the 500 random background sequences (**Fig. 3**).

To examine how different sequence contexts influence TF knockout effects, the motif analysis was repeated with motifs placed at the center of random sequences and putative enhancer sequences. Knockout effects were predicted using DREAM-RNN, and the distribution of effects across 250 random backgrounds and 1000 randomly selected putative enhancers was plotted to visualize how identical motif positions in different contexts can yield distinct knockout effects (**Fig. 4**). To compare motif knockout effects with a random control, random sequences of length 9–14 (range of the TF motif lengths used in this analysis) were generated and embedded at the center of the same backgrounds; knockouts predictions were performed in the same manner as above.

### *In silico* mutagenesis of variants to dissect strong positional variant effects

The variant exhibiting the largest change in both predicted and measured effects between different positions was selected for this analysis (**Fig. 5**). Changes in predicted expression were quantified as the sum of squared differences across positions *(50 bp upstream of center - variant placed at center)^2^ + (50 bp upstream of center - 50 bp downstream of center)^2^ + (50 bp downstream of center - variant placed at center)^2^*. Squaring ensured that changes in opposite directions did not cancel out, enabling identification of variants with the largest positional effects. To perform ISM, pretrained DREAM-RNN on K562 data from Gosai et al. was used to predict expression for the reference sequence and for sequences with each position mutated to each of the three other bases. Per base importance scores were computed as the average effect of mutating the reference allele to each of the three alternative bases (**Fig. 5a**).

For subsequent experiments, a potential interacting region (box B) was identified from the ISM. Box B was then mutated at its center, swapped with the variant region, reverse complemented, and tested with additional 5 bp or 10 bp insertions between motifs to assess how perturbations change variant effects and test for cooperativity (**Fig. 5b**). The exact sequences used for this analysis are available in Additional file 1: Table S1.

### Systematic interrogation of epistatic regulatory interactions

All Open Targets variants with PIP > 0.1 were analyzed, and their variant effects predicted when the variant was placed at the center of the sequence. To focus on strong-effect variants, those with predicted |log_2_FC| > 0.5 were selected. For each variant, ISM was performed for both the reference and alternate alleles, computing per-base importance scores as the average effect of mutating each base to the three alternative nucleotides.

To identify positions whose importance differs between the reference and alternate alleles (epistasis), and therefore potentially contribute to the variant effect, the change in importance scores between the two alleles was calculated. In this case, we compared the allele with higher predicted expression to that with lower predicted expression, ignoring reference and alternate status (since reference and alternate alleles are somewhat arbitrary). Finally, changes in importance scores were clustered by Euclidean distance to group similar effects together and the same clustering was applied to all plots (**Fig. 6**).

### Investigating selected variants

To determine whether variants exhibiting long-distance interacting regions were associated with known regulatory mechanisms, the sequences were searched against known database using BLAT and subsequently annotated with RepeatMasker to identify repetitive elements. For these variants, the aligned sequences were examined to assess alignment with bipartite Pol III promoter motifs. The interacting regions were compared against established consensus sequences for the internal promoter elements [34]:

- **Box A consensus:** TGGCTCACGCC
- **Box B consensus:** GWTCGAGAC (W = A or T)

Alignment to these consensus motifs (allowed up to 2 bp mismatch) was used to determine whether the interacting regions corresponded to box A and box B elements within *Alu* derived Pol III promoters and hence was used to characterize these variants into Pol III promoter variants.

## Abbreviations

bp: base pair
CRE: cis-regulatory element
eQTL: expression quantitative trait locus
FDR: false discovery rate
GTEx: Genotype-Tissue Expression
GWAS: genome-wide association study
ISM: in silico mutagenesis
KO: knockout
log2FC: log2 fold change
LSTM: long short-term memory
MPRA: massively parallel reporter assay
PIP: posterior inclusion probability
Pol III: RNA polymerase III
SINE: short interspersed nuclear element
SNV: single-nucleotide variant
TF: transcription factor
TFIIIB/TFIIIC: transcription factor IIIB/IIIC
TSS: transcription start site
UTR: untranslated region
WT: wild type

## Declarations

### Ethics approval and consent to participate

Not applicable. This study used only previously published, publicly available data.

### Consent for publication

Not applicable.

## Availability of data and materials

### Code Availability

All the scripts from the analysis in this paper and to recreate the figures can be found at https://github.com/de-Boer-Lab/position-mpra archived on Zenodo (https://zenodo.org/records/23488850) under the MIT license.

### Data Availability

All data used in this study are publicly available. Fine-mapped GWAS variants were obtained from the v2d_credset table of Open Targets Genetics (release 22.09) [44]. The K562 episomal MPRA data of Gosai et al. [8] used to train the models are available on Zenodo (https://doi.org/10.5281/zenodo.10698014) [46]. The K562 lentiMPRA data of Agarwal et al. [9] are available from the ENCODE portal under accession ENCSR382BVV (https://doi.org/10.17989/ENCSR382BVV; GEO accession GSE323223). The MPRA data that tested GTEx eQTL variants at three positions (Siraj et al. [22]) are available on Zenodo (https://doi.org/10.5281/zenodo.15297965).

## Authors’ contributions

S.I.A. performed the analyses, carried out the work described in this study and wrote the manuscript. C.G.D. supervised the project and R.T. provided guidance and feedback for analyzing and MPRA datasets. All authors read and approved the final manuscript

## Acknowledgements

This research was enabled in part by support provided by Advanced Research Computing at the University of British Columbia.

## Supplementary Information

Additional file 1 (PDF): Contains Supplementary Figures S1–S7 and Supplementary Table S1. Fig. S1: DREAM-RNN model trained on Gosai et al. K562 data has high predictive accuracy on held-out test set. Scatter plots comparing (a) predicted expression (x axis) versus actual expression (y axis) demonstrates strong correlation (Pearson’s correlation coefficient = 0.88), indicating robust model performance. (b) Scatterplots compare model-predicted effect (ref – alt) sizes (x-axis) to experimentally measured effects (ref – alt) (y-axis) for variants from Siraj et al. Separate panels show predictions evaluated with variants in different positions: all positions pooled, variant at the centre, and ±50 bp from centre. Each point represents a single variant. Supplementary Figure 2: Lentiviral MPRA, episomal MPRA, and K562 models trained on homology-leakage-filtered variants all demonstrate position-dependent variant effects across the probe sequence. Correlation of predicted effects (ref – alt) (colors) across variants decreases steadily as the distance (x and y axis) between predictions increases. (a) shows the correlation of all Open Target variants with PIP > 0.1 predicted using model trained on episomal MPRA (Gosai et al.), (b) shows the lentiMPRA (Agarwal et al.) trained model and (c) shows variants after homology leakage filtering, predicted using the model trained on Gosai et al. data. Fig. S3: Reproducibility between pseudo-replicate groups is high. Distribution of inter-replicate correlations, Pearson’s *r* (y axis) in MPRA-measured variant effects, calculated by randomly dividing the five experimental replicates into two pseudo-replicate groups through bootstrapping 10 times. The box spans the interquartile range (25th–75th percentile), the horizontal line indicates the median, and whiskers extend to the minimum and maximum values. Individual splits are shown as overlaid points. Fig. S4. Variant effects (ref–alt) vary by position for both the full set of fine-mapped variants with predicted |log_2_FC| > 0.5 at least one position (n = 33,319) and the subset remaining after homology leakage filtering (n = 1,302). (a) More variants have strong effects closer to the TSS. The % of variants with |log_2_FC| > 0.5 (y axis) at each position (x axis). (b) Similar positional activities are shared between variants. Variants (y axis) are clustered according to their scaled predicted log_2_FC (colors) for each position (x axis). Each variant is included twice, once in the reverse complement orientation. (c,d) The same analyses have been repeated for variants remaining after filtering out for homology leakage (n=1302), showing (c) the proportion of variants with |log_2_FC| > 0.5 across positions and (d) clustered positional activity profiles based on scaled predicted log_2_FC. Fig. S5: Both biochemical model and DREAM-RNN shows positional bias for yeast TFs. Positional activity scores (y axis), derived either from the biochemical model parameters (orange) or the DREAM-RNN TF knockout effect (blue), is displayed across various positions (x axis) upstream of the TSS where the motif center is placed. (a) and (b) refers to Mga1 in forward and reverse complemented orientation. Similarly, (c) and (d) refers to Skn7 and (e) and (f) refers to Ume6. For the DREAM-RNN predicted positional activity, the reported position (x axis) corresponds to the center of the motif relative to the TSS. In contrast, for the biochemical model, the position (x axis) refers to the filter position relative to the TSS, where the motif can occur at variable locations within the filter. This difference in how position is defined leads to the observed offset of a few bp between the two positional biases. Fig. S6: Distribution of Epistatic effects are strongest immediately adjacent to the variant and show a secondary peak at −65 bp. Each violin shows the distribution of epistasis scores (Δ ISM between the two alleles) (y-axis) at a given position (x-axis) relative to the variant (x=0), aggregated across all Open Target variants with |log_2_FC| > 0.5. Fig. S7: Variants with similar long-range interactions contain *Alu*-derived Pol III promoter elements. Nucleotides (colors) and position (x-axis) represent aligned coordinates post-alignment. Box A and B regions display high sequence homology to *Alu* Pol III promoter elements and sequences map to annotated *Alu* elements. Table S1: Systematic sequence perturbations for testing motif cooperative interactions. Summary of all sequence modifications performed on box A and box B. For each experimental perturbation, box A was tested in three distinct sequence contexts: variant placed at centre, 50 bp upstream of centre and 50 bp downstream of centre. Sequence modifications include motif mutation, position swapping, reverse complementation of box B and distance variations through 5 bp and 10 bp insertions. box A (orange) vs box B (blue) are highlighted in each sequence, and the alternate allele is highlighted in red (colors).

