## Supplemental figures and tables for "Position-dependent variant effects reveal importance of context in genomic regulation"

Supplementary Information

| **Perturbations** | **Reference Allele** | **Alternate Allele** |
| --- | --- | --- |
| box A and box B in original sequence | 50 bp upstream of centre:  GTGCCAATCCATCCCAAACATAATAGTTACAGATTGGCCGGGCGCGG**TGGCTCACGCC**TGTAATCCCAGCACTTTGGGAGGCCAAGGCGGATGGATCAGCTGAAGTCAGGA**GTTCGAGAC**CAGCCTGGCCAACATGGTGAAACCTTGTCTCTACTAAAAATACAAATATTAGCCAGGCGTCGTCGTGGGTGCCTGTGATC  variant placed at centre: TCTCCTCCAGGATTACTACTGTTAGTCTGTCTTTCCACCTCCAGTCTCTTGTGCCAATCCATCCCAAACATAATAGTTACAGATTGGCCGGGCGCGG**TGGCTCACGCC**TGTAATCCCAGCACTTTGGGAGGCCAAGGCGGATGGATCAGCTGAAGTCAGGA**GTTCGAGAC**CAGCCTGGCCAACATGGTGAAACCTTGTCT  50 bp downstream of centre: TTACATTTAAAATATTCCACATTCAGAGTTGTAGAGGCCTTAGACTATTATCTCCTCCAGGATTACTACTGTTAGTCTGTCTTTCCACCTCCAGTCTCTTGTGCCAATCCATCCCAAACATAATAGTTACAGATTGGCCGGGCGCGG**TGGCTCACGCC**TGTAATCCCAGCACTTTGGGAGGCCAAGGCGGATGGATCAGC | 50 bp upstream of centre: GTGCCAATCCATCCCAAACATAATAGTTACAGATTGGCCGGGCGCGG**TGCCTCACGCC**TGTAATCCCAGCACTTTGGGAGGCCAAGGCGGATGGATCAGCTGAAGTCAGGA**GTTCGAGAC**CAGCCTGGCCAACATGGTGAAACCTTGTCTCTACTAAAAATACAAATATTAGCCAGGCGTCGTCGTGGGTGCCTGTGATC  variant placed at centre: TCTCCTCCAGGATTACTACTGTTAGTCTGTCTTTCCACCTCCAGTCTCTTGTGCCAATCCATCCCAAACATAATAGTTACAGATTGGCCGGGCGCGG**TGCCTCACGCC**TGTAATCCCAGCACTTTGGGAGGCCAAGGCGGATGGATCAGCTGAAGTCAGGA**GTTCGAGAC**CAGCCTGGCCAACATGGTGAAACCTTGTCT  50 bp downstream of centre:  TTACATTTAAAATATTCCACATTCAGAGTTGTAGAGGCCTTAGACTATTATCTCCTCCAGGATTACTACTGTTAGTCTGTCTTTCCACCTCCAGTCTCTTGTGCCAATCCATCCCAAACATAATAGTTACAGATTGGCCGGGCGCGG**TGCCTCACGCC**TGTAATCCCAGCACTTTGGGAGGCCAAGGCGGATGGATCAGC |
| Mutate box B | 50 bp upstream of centre: GTGCCAATCCATCCCAAACATAATAGTTACAGATTGGCCGGGCGCGG**TGGCTCACGCC**TGTAATCCCAGCACTTTGGGAGGCCAAGGCGGATGGATCAGCTGAAGTCAGGA**GTTAGAGAC**CAGCCTGGCCAACATGGTGAAACCTTGTCTCTACTAAAAATACAAATATTAGCCAGGCGTCGTCGTGGGTGCCTGTGATC  variant placed at centre: TCTCCTCCAGGATTACTACTGTTAGTCTGTCTTTCCACCTCCAGTCTCTTGTGCCAATCCATCCCAAACATAATAGTTACAGATTGGCCGGGCGCGG**TGGCTCACGCC**TGTAATCCCAGCACTTTGGGAGGCCAAGGCGGATGGATCAGCTGAAGTCAGGA**GTTAGAGAC**CAGCCTGGCCAACATGGTGAAACCTTGTCT  50 bp downstream of centre:  TTACATTTAAAATATTCCACATTCAGAGTTGTAGAGGCCTTAGACTATTATCTCCTCCAGGATTACTACTGTTAGTCTGTCTTTCCACCTCCAGTCTCTTGTGCCAATCCATCCCAAACATAATAGTTACAGATTGGCCGGGCGCGG**TGGCTCACGCC**TGTAATCCCAGCACTTTGGGAGGCCAAGGCGGATGGATCAGC | 50 bp upstream of centre: GTGCCAATCCATCCCAAACATAATAGTTACAGATTGGCCGGGCGCGG**TGCCTCACGCC**TGTAATCCCAGCACTTTGGGAGGCCAAGGCGGATGGATCAGCTGAAGTCAGGA**GTTAGAGAC**CAGCCTGGCCAACATGGTGAAACCTTGTCTCTACTAAAAATACAAATATTAGCCAGGCGTCGTCGTGGGTGCCTGTGATC  variant placed at centre: TCTCCTCCAGGATTACTACTGTTAGTCTGTCTTTCCACCTCCAGTCTCTTGTGCCAATCCATCCCAAACATAATAGTTACAGATTGGCCGGGCGCGG**TGCCTCACGCC**TGTAATCCCAGCACTTTGGGAGGCCAAGGCGGATGGATCAGCTGAAGTCAGGA**GTTAGAGAC**CAGCCTGGCCAACATGGTGAAACCTTGTCT  50 bp downstream of centre:  TTACATTTAAAATATTCCACATTCAGAGTTGTAGAGGCCTTAGACTATTATCTCCTCCAGGATTACTACTGTTAGTCTGTCTTTCCACCTCCAGTCTCTTGTGCCAATCCATCCCAAACATAATAGTTACAGATTGGCCGGGCGCGG**TGCCTCACGCC**TGTAATCCCAGCACTTTGGGAGGCCAAGGCGGATGGATCAGC |
| Swap box A and box B | 50 bp upstream of centre:  GTGCCAATCCATCCCAAACATAATAGTTACAGATTGGCCGGGCGCGG**GTTCGAGAC**TGTAATCCCAGCACTTTGGGAGGCCAAGGCGGATGGATCAGCTGAAGTCAGGA**TGGCTCACGCC**  CAGCCTGGCCAACATGGTGAAACCTTGTCTCTACTAAAAATACAAATATTAGCCAGGCGTCGTCGTGGGTGCCTGTGATC  variant placed at centre: TCTCCTCCAGGATTACTACTGTTAGTCTGTCTTTCCACCTCCAGTCTCTTGTGCCAATCCATCCCAAACATAATAGTTACAGATTGGCCGGGCGCGG**GTTCGAGAC**TGTAATCCCAGCACTTTGGGAGGCCAAGGCGGATGGATCAGCTGAAGTCAGGA**TGGCTCACGCC**CAGCCTGGCCAACATGGTGAAACCTTGTCT  50 bp downstream of centre: TTACATTTAAAATATTCCACATTCAGAGTTGTAGAGGCCTTAGACTATTATCTCCTCCAGGATTACTACTGTTAGTCTGTCTTTCCACCTCCAGTCTCTTGTGCCAATCCATCCCAAACATAATAGTTACAGATTGGCCGGGCGCGG**TGGCTCACGCC**TGTAATCCCAGCACTTTGGGAGGCCAAGGCGGATGGATCAGC | 50 bp upstream of centre: GTGCCAATCCATCCCAAACATAATAGTTACAGATTGGCCGGGCGCGG**GTTCGAGAC**TGTAATCCCAGCACTTTGGGAGGCCAAGGCGGATGGATCAGCTGAAGTCAGGA**TGCCTCACGCC**  CAGCCTGGCCAACATGGTGAAACCTTGTCTCTACTAAAAATACAAATATTAGCCAGGCGTCGTCGTGGGTGCCTGTGATC  variant placed at centre: TCTCCTCCAGGATTACTACTGTTAGTCTGTCTTTCCACCTCCAGTCTCTTGTGCCAATCCATCCCAAACATAATAGTTACAGATTGGCCGGGCGCGG**GTTCGAGAC**TGTAATCCCAGCACTTTGGGAGGCCAAGGCGGATGGATCAGCTGAAGTCAGGA**TGCCTCACGCC**CAGCCTGGCCAACATGGTGAAACCTTGTCT  50 bp downstream of centre:  TTACATTTAAAATATTCCACATTCAGAGTTGTAGAGGCCTTAGACTATTATCTCCTCCAGGATTACTACTGTTAGTCTGTCTTTCCACCTCCAGTCTCTTGTGCCAATCCATCCCAAACATAATAGTTACAGATTGGCCGGGCGCGG**TGCCTCACGCC**TGTAATCCCAGCACTTTGGGAGGCCAAGGCGGATGGATCAGC |
| Revcomp box B | 50 bp upstream of centre:  GTGCCAATCCATCCCAAACATAATAGTTACAGATTGGCCGGGCGCGG**TGGCTCACGCC**TGTAATCCCAGCACTTTGGGAGGCCAAGGCGGATGGATCAGCTGAAGTCAGGA**GTCTCGAAC**CAGCCTGGCCAACATGGTGAAACCTTGTCTCTACTAAAAATACAAATATTAGCCAGGCGTCGTCGTGGGTGCCTGTGATC  variant placed at centre: TCTCCTCCAGGATTACTACTGTTAGTCTGTCTTTCCACCTCCAGTCTCTTGTGCCAATCCATCCCAAACATAATAGTTACAGATTGGCCGGGCGCGG**TGGCTCACGCC**TGTAATCCCAGCACTTTGGGAGGCCAAGGCGGATGGATCAGCTGAAGTCAGGA**GTCTCGAAC**CAGCCTGGCCAACATGGTGAAACCTTGTCT  50 bp downstream of centre: TTACATTTAAAATATTCCACATTCAGAGTTGTAGAGGCCTTAGACTATTATCTCCTCCAGGATTACTACTGTTAGTCTGTCTTTCCACCTCCAGTCTCTTGTGCCAATCCATCCCAAACATAATAGTTACAGATTGGCCGGGCGCGG**TGGCTCACGCC**TGTAATCCCAGCACTTTGGGAGGCCAAGGCGGATGGATCAGC | 50 bp upstream of centre: GTGCCAATCCATCCCAAACATAATAGTTACAGATTGGCCGGGCGCGG**TGCCTCACGCC**TGTAATCCCAGCACTTTGGGAGGCCAAGGCGGATGGATCAGCTGAAGTCAGGA**GTCTCGAAC**CAGCCTGGCCAACATGGTGAAACCTTGTCTCTACTAAAAATACAAATATTAGCCAGGCGTCGTCGTGGGTGCCTGTGATC  variant placed at centre: TCTCCTCCAGGATTACTACTGTTAGTCTGTCTTTCCACCTCCAGTCTCTTGTGCCAATCCATCCCAAACATAATAGTTACAGATTGGCCGGGCGCGG**TGCCTCACGCC**TGTAATCCCAGCACTTTGGGAGGCCAAGGCGGATGGATCAGCTGAAGTCAGGA**GTCTCGAAC**CAGCCTGGCCAACATGGTGAAACCTTGTCT  50 bp downstream of centre:  TTACATTTAAAATATTCCACATTCAGAGTTGTAGAGGCCTTAGACTATTATCTCCTCCAGGATTACTACTGTTAGTCTGTCTTTCCACCTCCAGTCTCTTGTGCCAATCCATCCCAAACATAATAGTTACAGATTGGCCGGGCGCGG**TGCCTCACGCC**TGTAATCCCAGCACTTTGGGAGGCCAAGGCGGATGGATCAGC |
| Insert 5bp in between box A and box B | 50 bp upstream of centre: GTGCCAATCCATCCCAAACATAATAGTTACAGATTGGCCGGGCGCGG**TGGCTCACGCC**TGTAATCCCAGCACTTTGGGAGGCCAAGGCGGATGGATCAGCTGAAGTCAGGAAGCTG**GTTCGAGAC**CAGCCTGGCCAACATGGTGAAACCTTGTCTCTACTAAAAATACAAATATTAGCCAGGCGTCGTCGTGGGTGCCTG  variant placed at centre : TCTCCTCCAGGATTACTACTGTTAGTCTGTCTTTCCACCTCCAGTCTCTTGTGCCAATCCATCCCAAACATAATAGTTACAGATTGGCCGGGCGCGG**TGGCTCACGCC**TGTAATCCCAGCACTTTGGGAGGCCAAGGCGGATGGATCAGCTGAAGTCAGGAAGCTG**GTTCGAGAC**CAGCCTGGCCAACATGGTGAAACCT  50 bp downstream of centre : TTACATTTAAAATATTCCACATTCAGAGTTGTAGAGGCCTTAGACTATTATCTCCTCCAGGATTACTACTGTTAGTCTGTCTTTCCACCTCCAGTCTCTTGTGCCAATCCATCCCAAACATAATAGTTACAGATTGGCCGGGCGCGG**TGGCTCACGCC**TGTAATCCCAGCACTTTGGGAGGCCAAGGCGGATGGATCAGC | 50 bp upstream of centre: GTGCCAATCCATCCCAAACATAATAGTTACAGATTGGCCGGGCGCGG**TGCCTCACGCC**TGTAATCCCAGCACTTTGGGAGGCCAAGGCGGATGGATCAGCTGAAGTCAGGAAGCTG**GTTCGAGAC**CAGCCTGGCCAACATGGTGAAACCTTGTCTCTACTAAAAATACAAATATTAGCCAGGCGTCGTCGTGGGTGCCTG  variant placed at centre : TCTCCTCCAGGATTACTACTGTTAGTCTGTCTTTCCACCTCCAGTCTCTTGTGCCAATCCATCCCAAACATAATAGTTACAGATTGGCCGGGCGCGG**TGCCTCACGCC**TGTAATCCCAGCACTTTGGGAGGCCAAGGCGGATGGATCAGCTGAAGTCAGGAAGCTG**GTTCGAGAC**CAGCCTGGCCAACATGGTGAAACCT  50 bp downstream of centre : TTACATTTAAAATATTCCACATTCAGAGTTGTAGAGGCCTTAGACTATTATCTCCTCCAGGATTACTACTGTTAGTCTGTCTTTCCACCTCCAGTCTCTTGTGCCAATCCATCCCAAACATAATAGTTACAGATTGGCCGGGCGCGG**TGCCTCACGCC**TGTAATCCCAGCACTTTGGGAGGCCAAGGCGGATGGATCAGC |
| Insert 10bp in between box A and box B | 50 bp upstream of centre: GTGCCAATCCATCCCAAACATAATAGTTACAGATTGGCCGGGCGCGG**TGGCTCACGCC**TGTAATCCCAGCACTTTGGGAGGCCAAGGCGGATGGATCAGCTGAAGTCAGGAAGCTGACGAT**GTTCGAGAC**CAGCCTGGCCAACATGGTGAAACCTTGTCTCTACTAAAAATACAAATATTAGCCAGGCGTCGTCGTGGGT  variant placed at centre : TCTCCTCCAGGATTACTACTGTTAGTCTGTCTTTCCACCTCCAGTCTCTTGTGCCAATCCATCCCAAACATAATAGTTACAGATTGGCCGGGCGCGG**TGGCTCACGCC**TGTAATCCCAGCACTTTGGGAGGCCAAGGCGGATGGATCAGCTGAAGTCAGGAAGCTGACGAT**GTTCGAGAC**CAGCCTGGCCAACATGGTGA  50 bp downstream of centre : TTACATTTAAAATATTCCACATTCAGAGTTGTAGAGGCCTTAGACTATTATCTCCTCCAGGATTACTACTGTTAGTCTGTCTTTCCACCTCCAGTCTCTTGTGCCAATCCATCCCAAACATAATAGTTACAGATTGGCCGGGCGCGG**TGGCTCACGCC**TGTAATCCCAGCACTTTGGGAGGCCAAGGCGGATGGATCAGC | 50 bp upstream of centre: GTGCCAATCCATCCCAAACATAATAGTTACAGATTGGCCGGGCGCGG**TGCCTCACGCC**TGTAATCCCAGCACTTTGGGAGGCCAAGGCGGATGGATCAGCTGAAGTCAGGAAGCTGACGAT**GTTCGAGAC**CAGCCTGGCCAACATGGTGAAACCTTGTCTCTACTAAAAATACAAATATTAGCCAGGCGTCGTCGTGGGT  variant placed at centre : TCTCCTCCAGGATTACTACTGTTAGTCTGTCTTTCCACCTCCAGTCTCTTGTGCCAATCCATCCCAAACATAATAGTTACAGATTGGCCGGGCGCGG**TGCCTCACGCC**TGTAATCCCAGCACTTTGGGAGGCCAAGGCGGATGGATCAGCTGAAGTCAGGAAGCTGACGAT**GTTCGAGAC**CAGCCTGGCCAACATGGTGA  50 bp downstream of centre : TTACATTTAAAATATTCCACATTCAGAGTTGTAGAGGCCTTAGACTATTATCTCCTCCAGGATTACTACTGTTAGTCTGTCTTTCCACCTCCAGTCTCTTGTGCCAATCCATCCCAAACATAATAGTTACAGATTGGCCGGGCGCGG**TGCCTCACGCC**TGTAATCCCAGCACTTTGGGAGGCCAAGGCGGATGGATCAGC |

**Table S1:** **Systematic sequence perturbations for testing motif cooperative interactions.** Summary of all sequence modifications performed on box A and box B. For each experimental perturbation, box A was tested in three distinct sequence contexts: variant placed at centre, 50 bp upstream of centre and 50 bp downstream of centre. Sequence modifications include motif mutation, position swapping, reverse complementation of box B and distance variations through 5 bp and 10 bp insertions. box A (orange) vs box B (blue) are highlighted in each sequence, and the alternate allele is highlighted in red (colors).

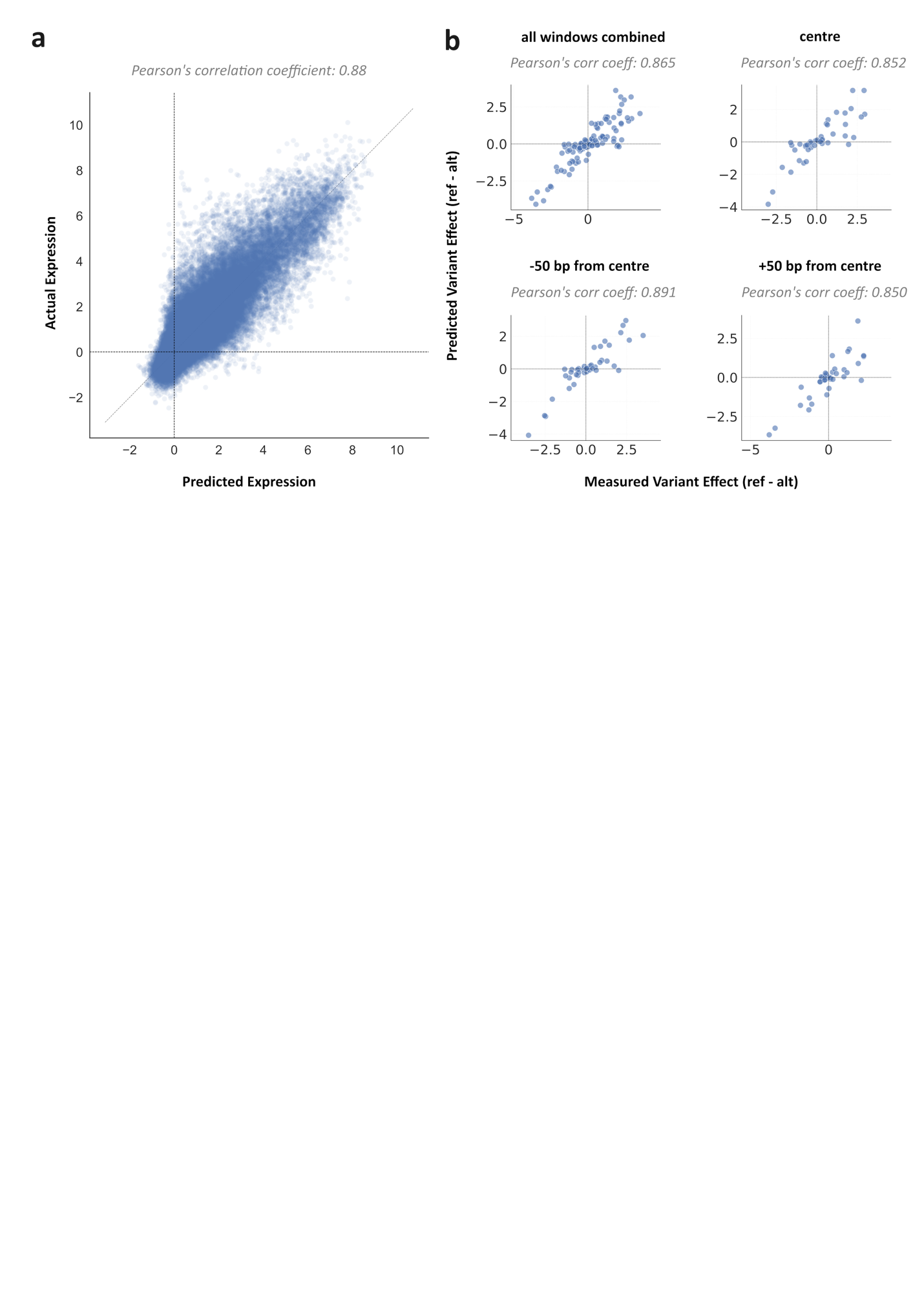
**Fig. S1: DREAM-RNN model trained on Gosai et al. K562 data has high predictive accuracy on held-out test set.** Scatter plots comparing (a) predicted expression (x axis) versus actual expression (y axis) demonstrates strong correlation (Pearson’s correlation coefficient = 0.88), indicating robust model performance. (b) Scatterplots compare model-predicted effect (ref – alt) sizes (x-axis) to experimentally measured effects (ref – alt) (y-axis) for variants from Siraj et al. Separate panels show predictions evaluated with variants in different positions: all positions pooled, variant at the centre, and ±50 bp from centre. Each point represents a single variant.

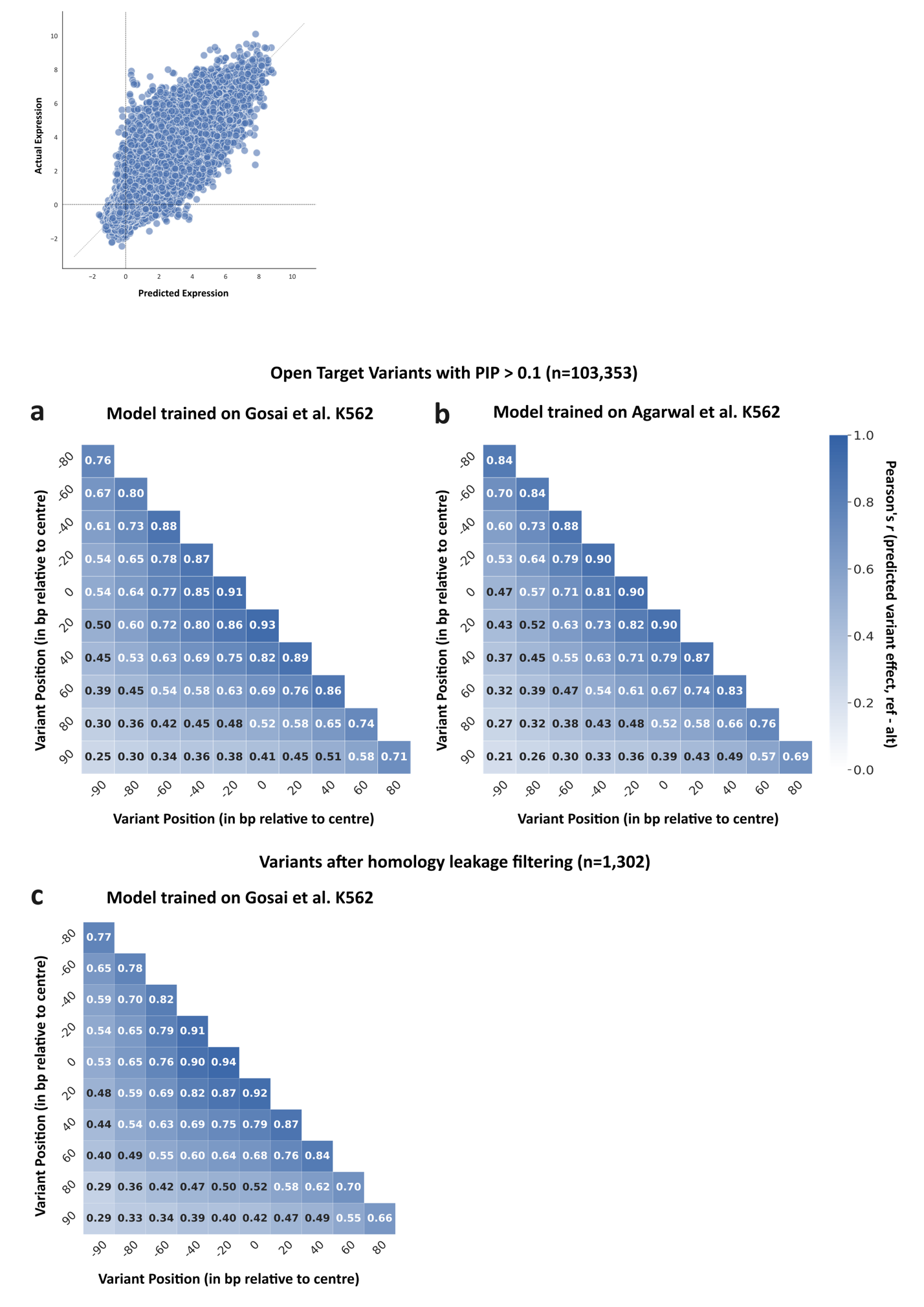

**Supplementary Figure 2: Lentiviral MPRA, episomal MPRA, and K562 models trained on homology-leakage-filtered variants all demonstrate position-dependent variant effects across the probe sequence.** Correlation of predicted effects (ref – alt) (colors) across variants **decreases steadily as the distance (x and y axis) between predictions increases. (a) shows the correlation of all Open Target variants with PIP > 0.1 predicted using model trained on episomal MPRA (Gosai et al.), (b) shows the lentiMPRA (Agarwal et al.) trained model and** (c) shows variants after homology leakage filtering, predicted using the model trained on Gosai et al. data.

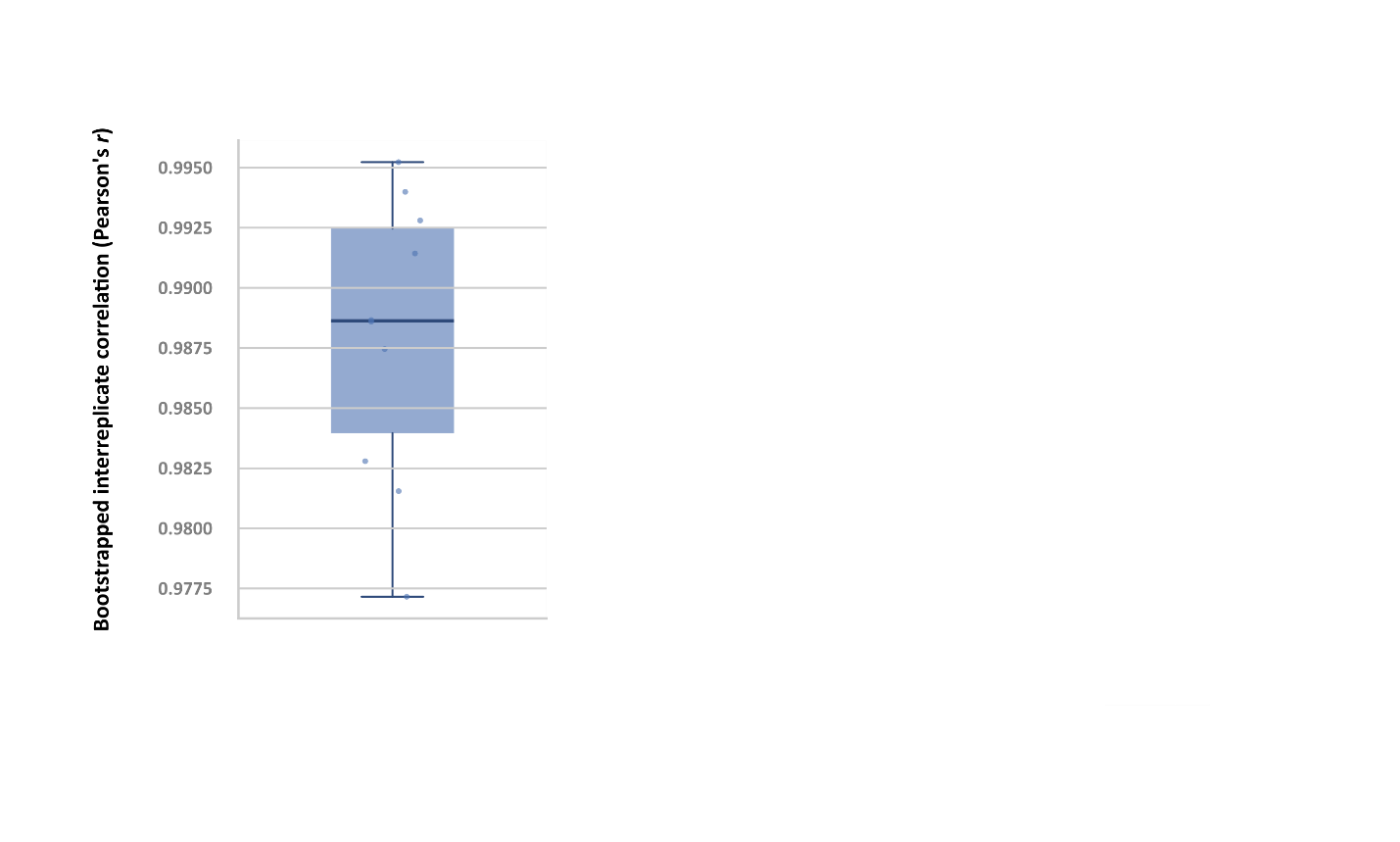

**Fig. S3:** **Reproducibility between pseudo-replicate groups is high.** Distribution of inter-replicate correlations, Pearson’s *r* (y axis) in MPRA-measured variant effects, calculated by randomly dividing the five experimental replicates into two pseudo-replicate groups through bootstrapping 10 times. The box spans the interquartile range (25th–75th percentile), the horizontal line indicates the median, and whiskers extend to the minimum and maximum values. Individual splits are shown as overlaid points.

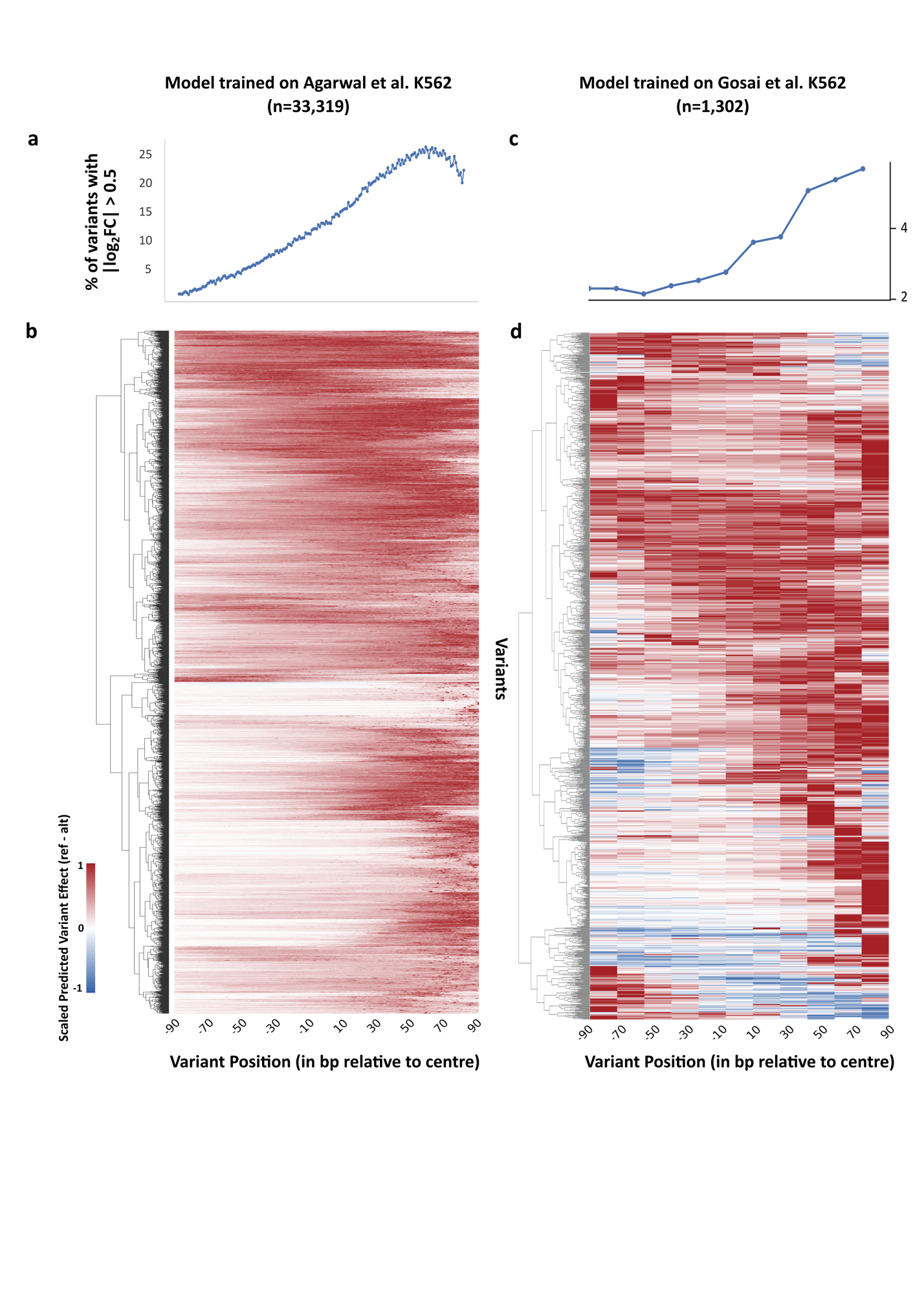

**Fig. S4: Variant effects (ref–alt) vary by position for both the full set of fine-mapped variants with predicted |log_2_FC| > 0.5 at least one position (n = 33,319) and the subset remaining after homology leakage filtering (n = 1,302).** **(a)** More variants have strong effects closer to the TSS. The % of variants with |log_2_FC| > 0.5 (y axis) at each position (x axis). **(b)** Similar positional activities are shared between variants. Variants (y axis) are clustered according to their scaled predicted log_2_FC (colors) for each position (x axis). Each variant is included twice, once in the reverse complement orientation. **(c,d)** The same analyses have been repeated for variants remaining after filtering out for homology leakage (n=1302), showing **(c)** the proportion of variants with |log_2_FC| > 0.5 across positions and **(d)** clustered positional activity profiles based on scaled predicted log_2_FC.

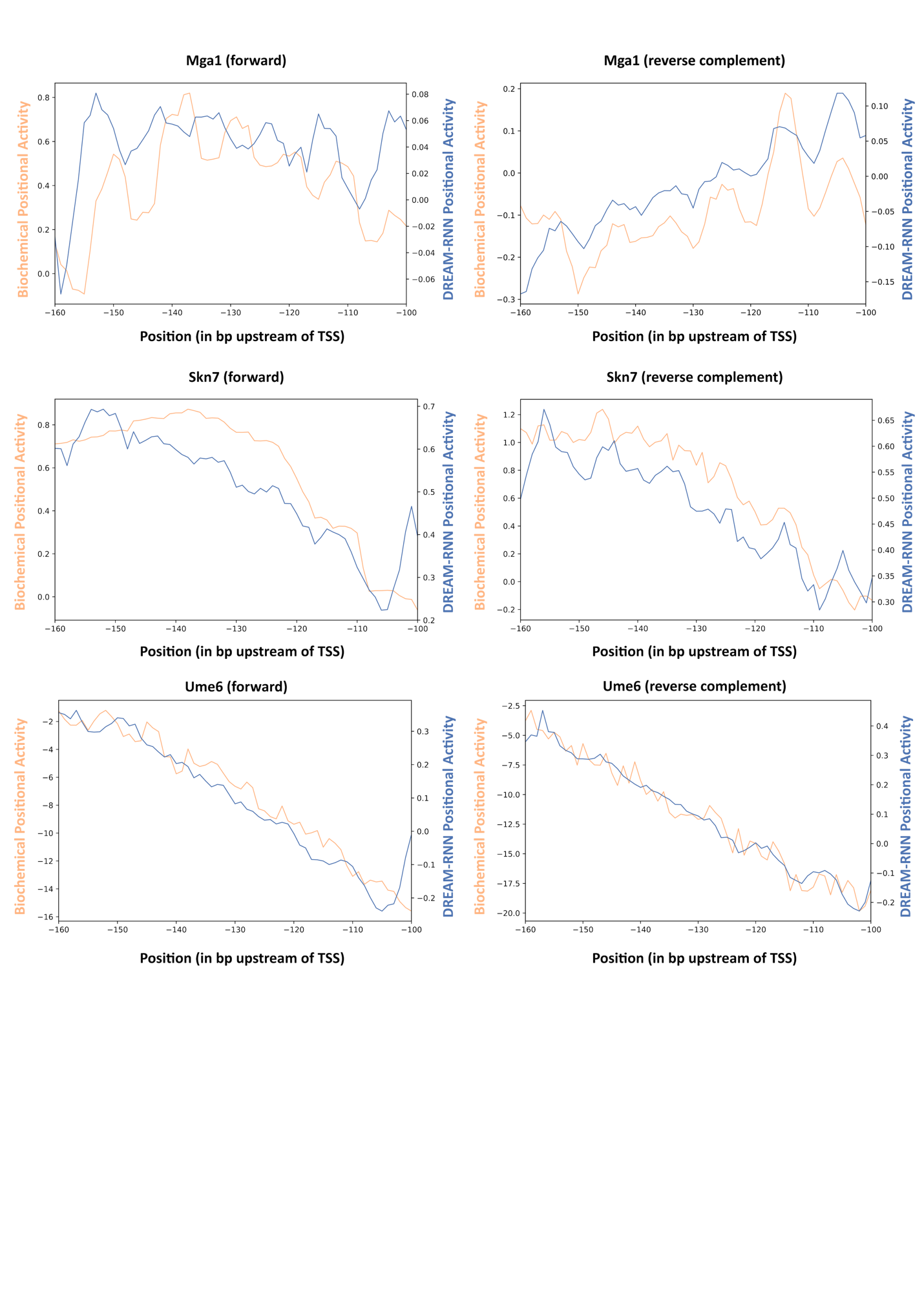

**Fig. S5: Both biochemical model and DREAM-RNN shows positional bias for yeast TFs.** Positional activity scores (y axis), derived either from the biochemical model parameters (orange) or the DREAM-RNN TF knockout effect (blue), is displayed across various positions (x axis) upstream of the TSS where the motif center is placed. (a) and (b) refers to Mga1 in forward and reverse complemented orientation. Similarly, (c) and (d) refers to Skn7 and (e) and (f) refers to Ume6. For the DREAM-RNN predicted positional activity, the reported position (x axis) corresponds to the **center of the motif relative to the TSS**. In contrast, for the biochemical model, the position (x axis) refers to the **filter position relative to the TSS**, where the motif can occur at variable locations within the filter. This difference in how position is defined leads to the observed offset of a few bp between the two positional biases.
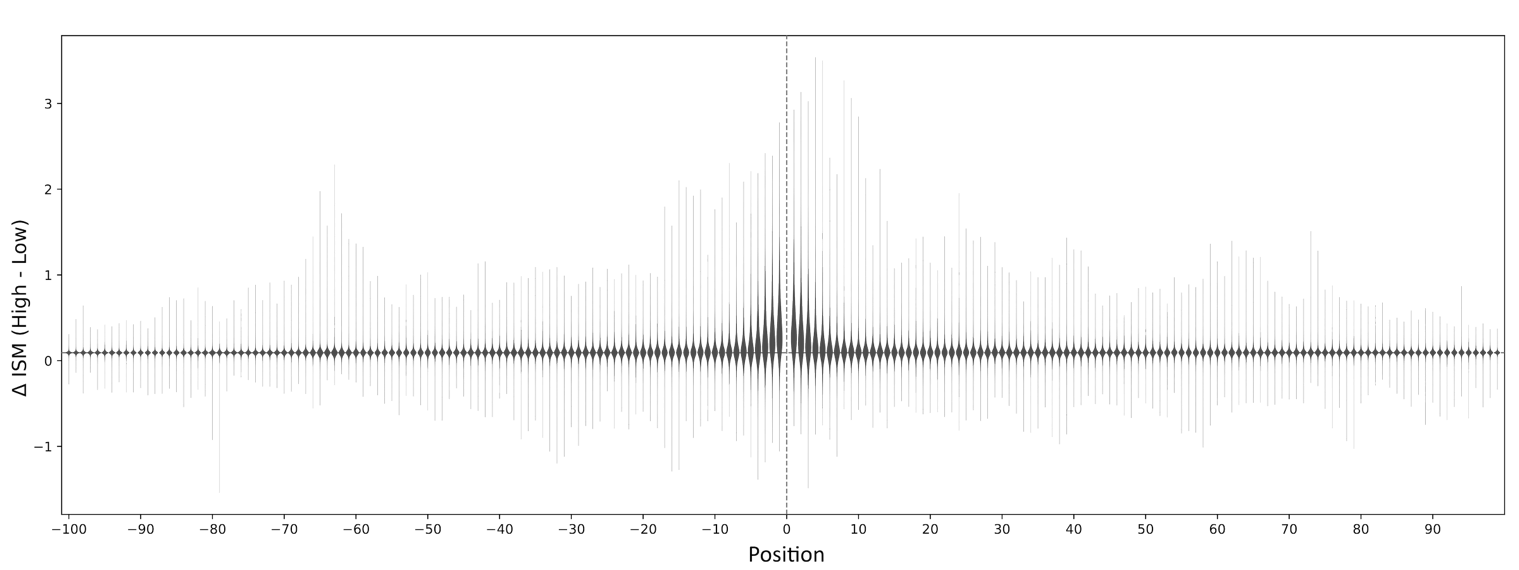

**Fig. S6: Distribution of Epistatic effects are strongest immediately adjacent to the variant and show a secondary peak at −65 bp.** Each violin shows the distribution of epistasis scores (Δ ISM between the two alleles) (y-axis) at a given position (x-axis) relative to the variant (x=0), aggregated across all Open Target variants with |log_2_FC| > 0.5.

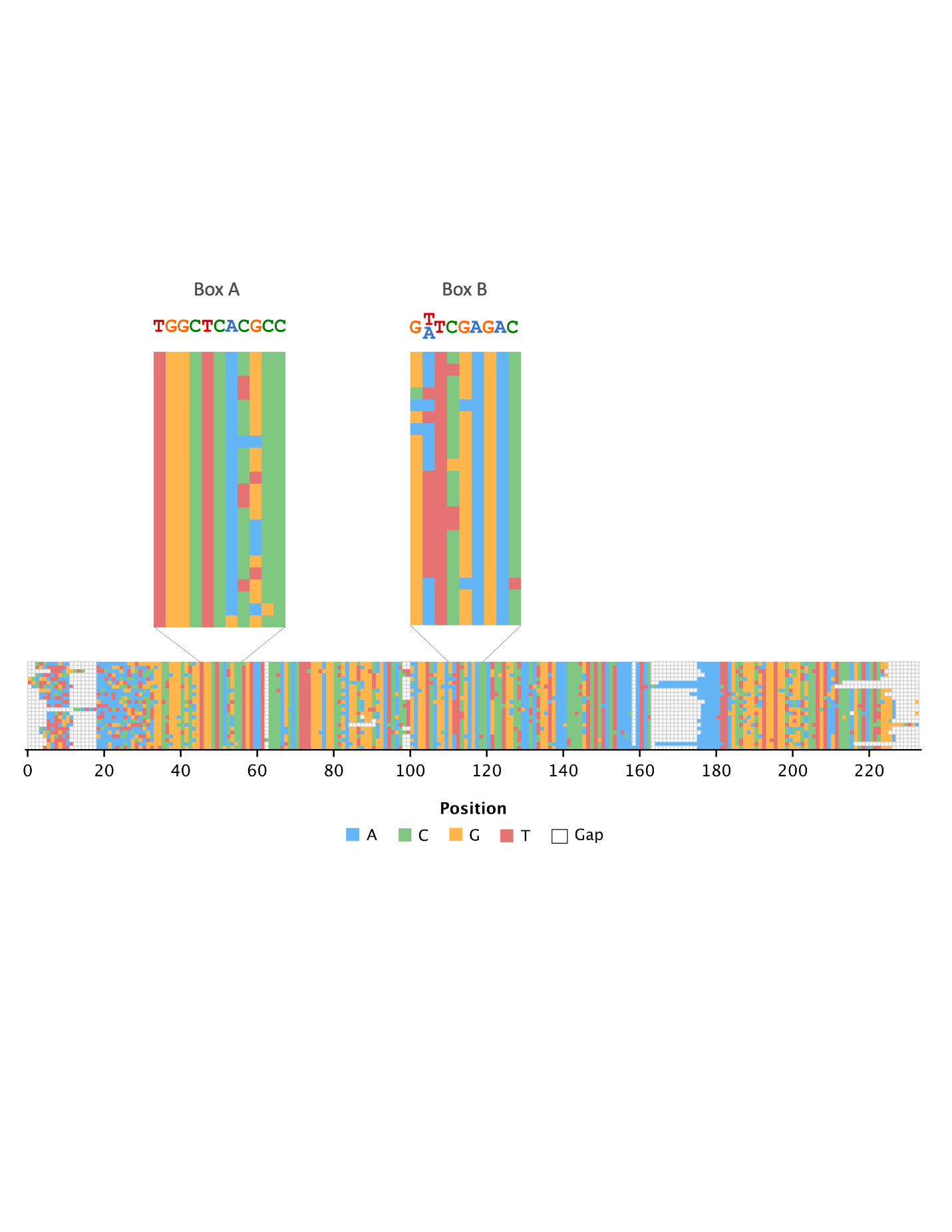

**Fig. S7:** **Variants with similar long-range interactions contain *Alu*-derived Pol III promoter elements.** Nucleotides (colors) and position (x-axis) represent aligned coordinates post-alignment. Box A and B regions display high sequence homology to *Alu* Pol III promoter elements and sequences map to annotated *Alu* elements.
